# Murine Alveolar Macrophages Rapidly Accumulate Intranasally Administered SARS-CoV-2 Spike Protein leading to Neutrophil Recruitment and Damage

**DOI:** 10.1101/2023.03.13.532446

**Authors:** Chung Park, Il-Young Hwang, Serena Li-Sue Yan, Sinmanus Vimonpatranon, Danlan Wei, Don Van Ryk, Alexandre Girard, Claudia Cicala, James Arthos, John H. Kehrl

**Author notes:** Corresponding author, To whom correspondence should be addressed: John H. Kehrl, Laboratory of Immunoregulation, National Institute of Allergy and Infectious Diseases, National Institutes of Health, Bldg. 10, Room 6A01, 10 Center Dr. MSC 1876, Bethesda, Maryland 20892, United States of America.

## Abstract

The trimeric SARS-CoV-2 Spike protein mediates viral attachment facilitating cell entry. Most COVID-19 vaccines direct mammalian cells to express the Spike protein or deliver it directly via inoculation to engender a protective immune response. The trafficking and cellular tropism of the Spike protein *in vivo* and its impact on immune cells remains incompletely elucidated. In this study we inoculated mice intranasally, intravenously, and subcutaneously with fluorescently labeled recombinant SARS-CoV-2 Spike protein. Using flow cytometry and imaging techniques we analyzed its localization, immune cell tropism, and acute functional impact. Intranasal administration led to rapid lung alveolar macrophage uptake, pulmonary vascular leakage, and neutrophil recruitment and damage. When injected near the inguinal lymph node medullary, but not subcapsular macrophages, captured the protein, while scrotal injection recruited and fragmented neutrophils. Wide-spread endothelial and liver Kupffer cell uptake followed intravenous administration. Human peripheral blood cells B cells, neutrophils, monocytes, and myeloid dendritic cells all efficiently bound Spike protein. Exposure to the Spike protein enhanced neutrophil NETosis and augmented human macrophage TNF-α and IL-6 production. Human and murine immune cells employed C-type lectin receptors and Siglecs to help capture the Spike protein. This study highlights the potential toxicity of the SARS-CoV-2 Spike protein for mammalian cells and illustrates the central role for alveolar macrophage in pathogenic protein uptake.

## Introduction

In 2019 a new coronavirus (SARS-CoV-2) was identified as the cause of an epidemic outbreak of an acute respiratory syndrome in Wuhan, China. SARS-CoV-2 used the same cell entry receptor—angiotensin converting enzyme II (ACE2)—as did SARS-CoV-1 (1). The SARS-CoV-2 Spike protein mediates cell entry and is a single-pass transmembrane proteins that forms homotrimers. It has a large N-terminal ectodomain exposed to the exterior, a transmembrane helix, and a short C-terminal tail located within the virus. Each Spike monomer contains two regions termed S1 and S2. In the assembled trimer the S1 regions contain the receptor binding domain while the S2 regions forms a flexible stalk, which mediates membrane fusion between the viral envelope and the host cell membrane (2–4). The S1 region is divided into a N-terminal domain and C-terminal domain, the latter interacts with the target cell ACE2 (5–11). Like other Coronavirus spike proteins SARS-CoV-2 Spike protein is heavily N-linked glycosylated (12, 13) and blocking N- and O-glycans dramatically reduced viral entry (14). The SARS-CoV-2 Spike proteins is activated by host cell proteases that cleave the protein at the S1-S2 boundary and subsequently at the S2’ site (15–17). Because Spike proteins are located on the surface of the virus, they are a major antigen targeted by the host immune system (18).

During natural infection host immune cells encounter Spike proteins via several different avenues. First, by direct contact with Spike protein bearing viral particles released from infected cells. Second, although the SARS-CoV-2 virions assemble in intracellular compartments of infected cells, unincorporated Spike proteins can reach the plasma membrane. Infected cells expressing Spike proteins may bind to cellular receptors present on resident or recruited immune cells. Third, extracellular vesicles (EVs) released by virally infected cells can contain Spike proteins. Mass spectrometry and nanoscale flow cytometry demonstrated SARS-CoV-2 Spike protein incorporation into EVs (19). Fourth, Spike proteins are preprocessed during viral assembly utilizing the furin protease cleavage site between the S1 and S2 subunits (17). When the Spike protein adopts a fusion conformation the ACE2 receptor-binding domain separates from the membrane-bound S2 subunit. Soluble S1 subunits shed from infected cells or from the virions *in vivo* may bind to other cells via ACE2 or other binding partners. Intriguingly 60% of the plasma samples from patients with post-acute sequelae of coronavirus disease 2019 (PASC) had detectable levels of Spike protein using an ultrasensitive antigen capture assay (20). Prolonged exposure to Spike protein has been suggested to be responsible for Long-COVID syndrome (21). Humans also encounter SARS-CoV-2 Spike protein via vaccination (22). Serious adverse events following vaccination are rare but include vaccine-induced immune thrombocytopenia and thrombosis; myocarditis and pericarditis; and a variety of autoimmune illnesses (23). Due to waning effectiveness humans require repeated immunizations to maintain immunity and protection against potentially severe disease raising some concerns about the impact of repeated exposure to the SARS-CoV-2 Spike proteins.

To better understand the localization and trafficking of the SARS-CoV-2 Spike protein following administration and perhaps during natural infection we prepared a recombinant SARS-CoV-2 Spike ectodomain stabilized in a prefusion conformation (24). This variant (S-2P) contained two consecutive proline substitutions in the S2 subunit. This double-proline substitution (SARS-CoV-2 S-2P) has allowed the rapid determination of high-resolution cryo-EM structures. Fluorescently labeled SARS-CoV-2 Spike ectodomain, a D614G variant (25–27), a high mannose (14), and de-glycosylated version were injected into mice to characterize uptake and identify human mononuclear cells that bound these proteins, and to perform functional studies. In some instances, we used viral like particles (VLPs) expressing the full-length Spike protein. Our results identified the *in vivo* cellular tropism of the SARS-CoV-2 Spike proteins, delineated their human mononuclear cell targets, provided insights into their functional effects, thereby, helping afford a basis for understanding their impact on humans.

## Results

### Preparation of SARS-CoV-2 Spike proteins and VLPs expressing them

We purified a stabilized exodomain of the original SARS-CoV-2 Spike protein (24) using media conditioned by transfected CHO-S or HEK293F cells. Because of higher yields most experiments used CHO-S derived protein. The strategy for producing the recombinant protein is shown (**Figure 1-figure supplement 1**). Based on its mobility on size exclusion chromatography the Spike proteins spontaneously formed trimers. Kifunensine treated cultures were used to generate a high mannose version, and PNGase F treated protein generated an N-glycan deficient version, and they will be referred to as such. The PNGase F treatment resulted in two distinct chromatography peaks that exhibited slightly different mobilities on SDS-PAGE. As discussed below we predominately used the preparation from fractions 9-11. The D614G mutant of the original Wuhan SARS-CoV-2 Spike protein was also purified. Each of the recombinant proteins was subjected to two rounds of Triton X-114 extraction to remove any residual lipopolysaccharide (28). The N-termini of the recombinant proteins were labeled with Alexa Fluor 488. In some experiments we used viral like particles (VLPs) that expressed the original Wuhan Spike protein. We produced the VLPs using HEK293T cells along with a plasmid encoding the human immunodeficiency GAG protein fused to green fluorescent protein (GFP) and a full-length SARS-CoV-2 Spike protein expression vector. Mice received the Spike protein preparations or the SARS-CoV-2 VLPs by intranasal instillation, while only the Spike proteins were injected intravenously or subcutaneously.

**Figure 1.**
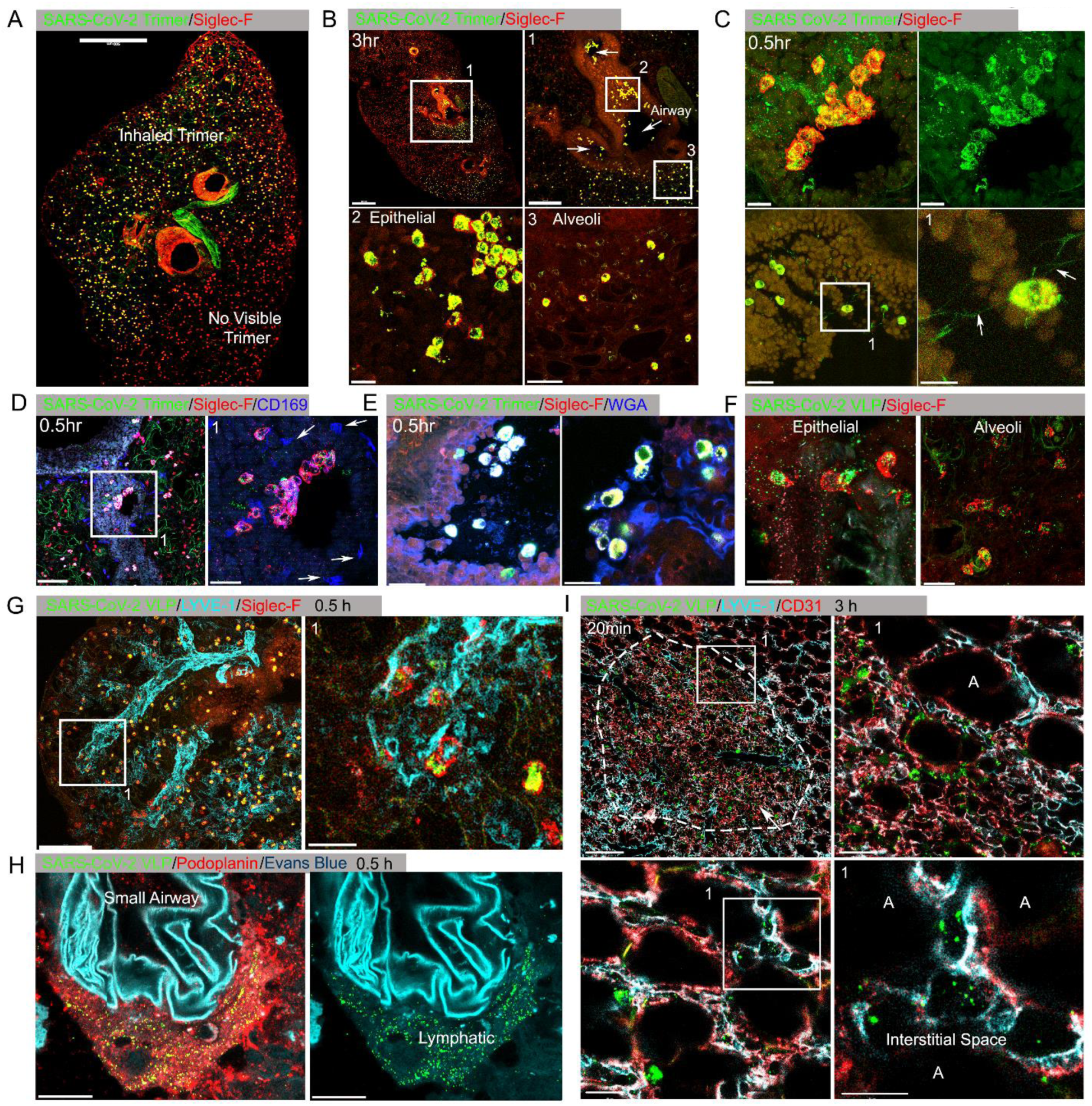
Lung images following intranasal administered SARS-CoV-2 Spike protein. **(A)** Confocal micrograph of a lung section 18 hr post SARS-CoV-2 Spike protein (Trimer) instillation. Alveolar macrophages (AMs) visualized with Siglec-F antibody. Alexa Fluor 488 conjugated recombinant SARS-CoV-2 Spike protein (3 μg in 50 μl of saline) was inoculated by nasal instillation. Spike-reached lung region (Instilled Trimer) and non-reached area (No visible Trimer) noted on the micrograph. Scale bar, 500 µm. **(B)** Confocal micrographs of lung collected at 3 hr post Spike protein instillation. AMs in the large airway and alveoli (upper left) area is shown. ROI-1 (box1) (upper right) is enlarged, and arrows indicate large airways. ROI-2 (Box 2) (lower left) shows AMs bearing SARS-CoV-2 Spike protein on airway epithelial cells. ROI-3 (Box 3) (lower left) shows Spike protein bearing AMs in alveoli. Scale bars, 500, 200, 20, and 50 μm. (**C)** Confocal micrographs of lung collected at 0.5 hr post SARS-CoV-2 Spike protein instillation. AMs were visualized with Siglec-F antibody (upper left & right). AMs connected to each other (lower left) via tunneling nanotubes (arrows) (lower right). Scale bars, 20, 20, 30, and 10 μm. **(D & E)** Confocal micrographs of lung collected at 0.5 hr post Spike instillation. SARS-CoV-2 Spike protein (green) on airway epithelium and CD169^+^ macrophages (D). Arrows indicate CD169^+^ macrophages (right). Scale bars, 100 and 30 μm. AMs with SARS-CoV-2 Spike protein and Mucin (Alexa Fluor 647 conjugated wheat germ agglutin, WGA) on airway epithelium (E). Scale bars, 30 and 20 μm. **(F)** Confocal micrographs of lung collected at 2 hr post SARS-CoV-2 Spike protein incorporated VLP instillation (green). AMs (red, SiglecF) on epithelium (left) and in alveoli (right). Scale bars, 20 μm. **(G)** Confocal micrographs of lung collected at 0.5 hr post SARS-CoV-2 Spike VLP (green) instillation. AMs (red, Siglec-F antibody) and lung vasculatures visualized LYVE-1 (cyan) (top, left). LYVE-1^+^ vasculature associated AMs bearing VLPs is highlighted (box) and enlarged (top, right). Scale bars, 100 and 25 μm. **(H)** A confocal micrograph shows a lung lymphatic vasculature visualized with Podoplanin antibody. Fifty microliters of a mixture of Evans blue (cyan) (5 μg) and Spike bearing VLPs (green) (0.5 million counts) were applied to the mouse nose. VLPs in lymphatics associated with a small airway highlighted (right). Scale bars, 20 μm. **(I)** Confocal micrographs of lung collected 20 min post Spike incorporated VLP (green) instillation. Lung vasculatures visualized CD31 (red) and LYVE-1 (cyan). Damaged lung tissue is indicated (dotted line) (upper left). Border of damaged tissue and intact alveoli (box) is enlarged (upper right). A lymphatic structure was stained by LYVE-1 in intact alveolus is highlighted (lower left). An image of VLPs in LYVE-1^+^ lymphatic portal in alveoli (box) is enlarged (lower right). “A” indicates alveolus. Scale bars, 100, 20, 25, and 10 μm.

### Instillation of SARS-CoV-2 Spike proteins and Spike protein expressing VLPs affects the lung cellular composition and architecture

At various time points after Spike protein or VLP nasal instillation lungs from mice were processed for confocal microscopy. A Siglec-F antibody identified alveolar macrophages (AMs) while CD169 (Siglec-1) immunostaining delineated a subset of large airway associated macrophages (29). Interstitial and inflammatory macrophages do not express Siglec-F. At 3 hours post instillation low magnification images of mouse lung sections revealed Spike protein (Trimer) uptake by Siglec-F positive cells. Due to the uneven distribution of the instilled Spike protein the Siglec-F positive cells in the lower portion of the image lack signal (**Figure 1A**, **Figure 1-figure supplement 2A**). The Siglec-F positive macrophages located near the bronchial epithelium and in the nearby alveoli initially accumulated the labeled protein **(Figure 1B)**. Even at the 30-minute time point AMs had already accumulated it **(Figure 1C)**. Most of the macrophages that acquired the Spike proteins were Siglec-F positive, lacked CD169 **(Figure 1D)**, but were wheat germ agglutinin positive, a mucin marker **(Figure 1E)**. Repeating the experiment but substituting the Spike protein with VLPs expressing the SARS-CoV-2 Spike protein revealed a similar pattern of uptake by AMs, but with greater neutrophil recruitment and granulation compared to delta Env VLPs. **(Figure 1F**, **Figure 1-figure supplement 2B, C)**. Immunostaining with LYVE-1 and CD31 to identify the lymphatic and blood vessel endothelium, respectively, **(Figure 1G)** or with Podoplanin another lymphatic endothelial cell marker **(Figure 1H**, **Figure 1-figure supplement 2D)** showed the rapid accumulation of the Spike protein bearing VLPs in small and large lymphatic vessels. Within 3 hours of administration of either the VLPs **(Figure 1I)** or the Spike protein (data not shown) areas of alveolar collapse and lung damage were evident.

To assess the cellular response to Spike protein instillation and to delineate the cells that had acquired Spike protein *in vivo* we collected mouse lungs 18 hours post exposure to different SARS-CoV-2 Spike protein preparations. Saline instillation served as a control. Lung fragments were digested, and then pushed through a 40-micron cell strainer. Analysis of the collected cells by flow cytometry revealed that the SARS-CoV-2 Spike protein increased the number of neutrophils, monocytes, and dendritic cells in the lung, but did not affect lymphocyte or eosinophil numbers **(Figure 2A)**. Due to the uneven distribution of the instilled protein and because of their rapid turnover the neutrophil numbers likely underestimate the local neutrophil recruitment. As the imaging experiments indicated AMs most avidly acquired the Spike protein, some interstitial macrophages also retained it, while only a low percentage of the lung neutrophils and dendritic cells were positive following isolation **(Figure 2B)**. Instillation of the glycan deficient Spike protein reduced the monocyte cellular infiltrate and decreased the % of alveolar macrophages that retained the protein. The high mannose Spike protein slightly increased the numbers of neutrophil and macrophages in the lung despite a lower uptake by alveolar and interstitial macrophages. Finally, the D614G mutation reduced the cellular infiltrate compared to the unmutated protein and had a slightly different cell binding profile as a greater percentage of monocytes, eosinophils, and dendritic cells retained it **(Figure 2A, B)**. We also tested whether the addition of human ACE2 affected the cellular uptake of the Spike protein following intranasal administration by comparing wild type and the human ACE2 transgenic mice. While we saw little difference by imaging, we did note some minor changes in the cellular uptake pattern, most notably an increase uptake by AM and decrease by interstitial macrophages **(Figure 2-figure supplement 1)**

**Figure 2.**
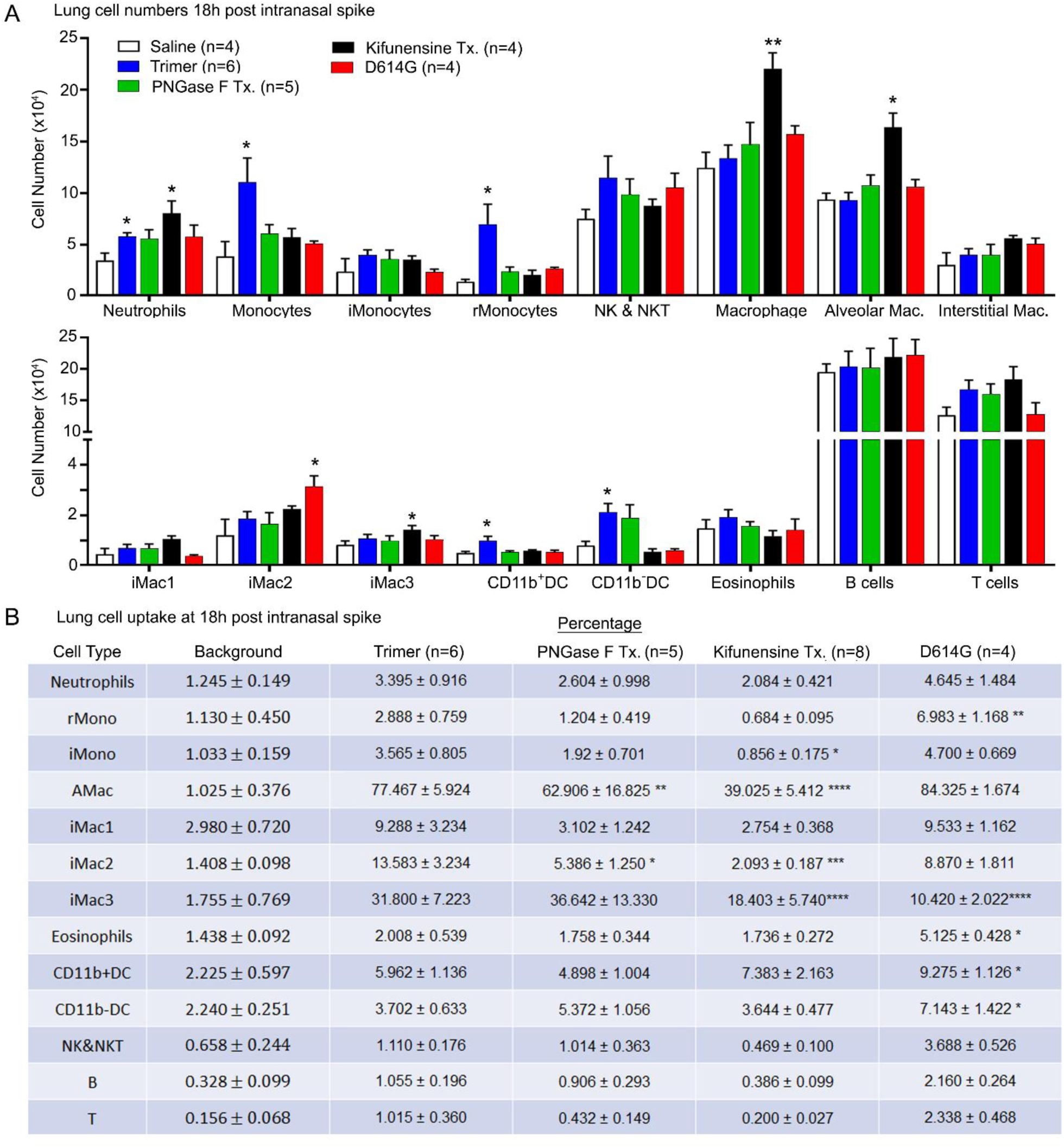
Lung leukocyte profile following intranasal administration of SARS-CoV-2 Spike proteins. **(A)** Leukocyte numbers in isolated lung tissue. Eighteen hr post intranasal administration of indicated Spike proteins lung tissue collected, processed, and leukocyte cell numbers determined by flow cytometry. Numbers of mice analyzed are indicated. Data mean +/− SEM, *p < 0.05; **p < 0.005. **(B)** SARS-CoV-2 Spike protein uptake by lung leukocytes 18 hr following intranasal inoculation. Flow cytometry results from indicated mice. Values significantly different from the WT SARS-CoV-2 protein (Trimer) are indicated, *p < 0.05; **p < 0.01; ***p < 0.005. ****p<0.001.

### Altered lung vascular permeability, neutrophil recruitment, and lung damage 3 hours post instillation of the Spike protein

The increase lung leukocytes following the Spike protein nasal instillation suggested that it may have altered the vascular permeability of the lung. To assess whether vasculature permeability changes had occurred, we intravenously injected Evans blue dye, which in the absence of a permeability defect remains confined to the vasculature (30). Prior to injecting the Evans blue, we instilled in the nasal cavity the SARS-CoV-2 Spike protein, the S1 subunit of the HCoV-HKU1 Spike protein, or a saline control. Inspection of the lungs from the SARS-CoV-2 spike protein treated mice revealed a strong increase in Evan blue staining while the lungs from the S1 subunit HCoV-HKU1 treated mice had a minimal increase. Quantifying the Evans blue dye in collected lungs and liver confirmed the increase in vasculature permeability in the lungs of the SARS-CoV-2 Spike protein treated mice **(Figure 3A).** Next, we compared the S1 subunit of SARS-CoV-2 Spike protein to the HCoV-HKU1 S1 subunit. The S1 subunit preparations were purified from transfected HEK 293 cells. The results of administrating the S1 subunit of HCoV-HKU1 Spike protein did not differ from the saline control while the S1 subunit from SARS-CoV-2 like the SARS-CoV-2 trimer increased the lung vascular permeability to Evans blue (**Figure 3-figure supplement 1A)**. We also tested whether the presence of hACE2 affected the response by using K18-hACE2 transgenic mice as the recipients. The presence of the K18-hACE2 transgene slightly increased the lung vasculature permeability upon SARS-CoV-2 trimer instillation **(Figure 3-figure supplement 1B).** Consistent with the increase in lung vascular permeability 3 hours post SARS-CoV-2 Spike protein instillation, Siglec-F positive lung macrophages bearing the Spike protein were surrounded by neutrophils and neutrophil fragments **(Figure 3B, C)**.

**Figure 3.**
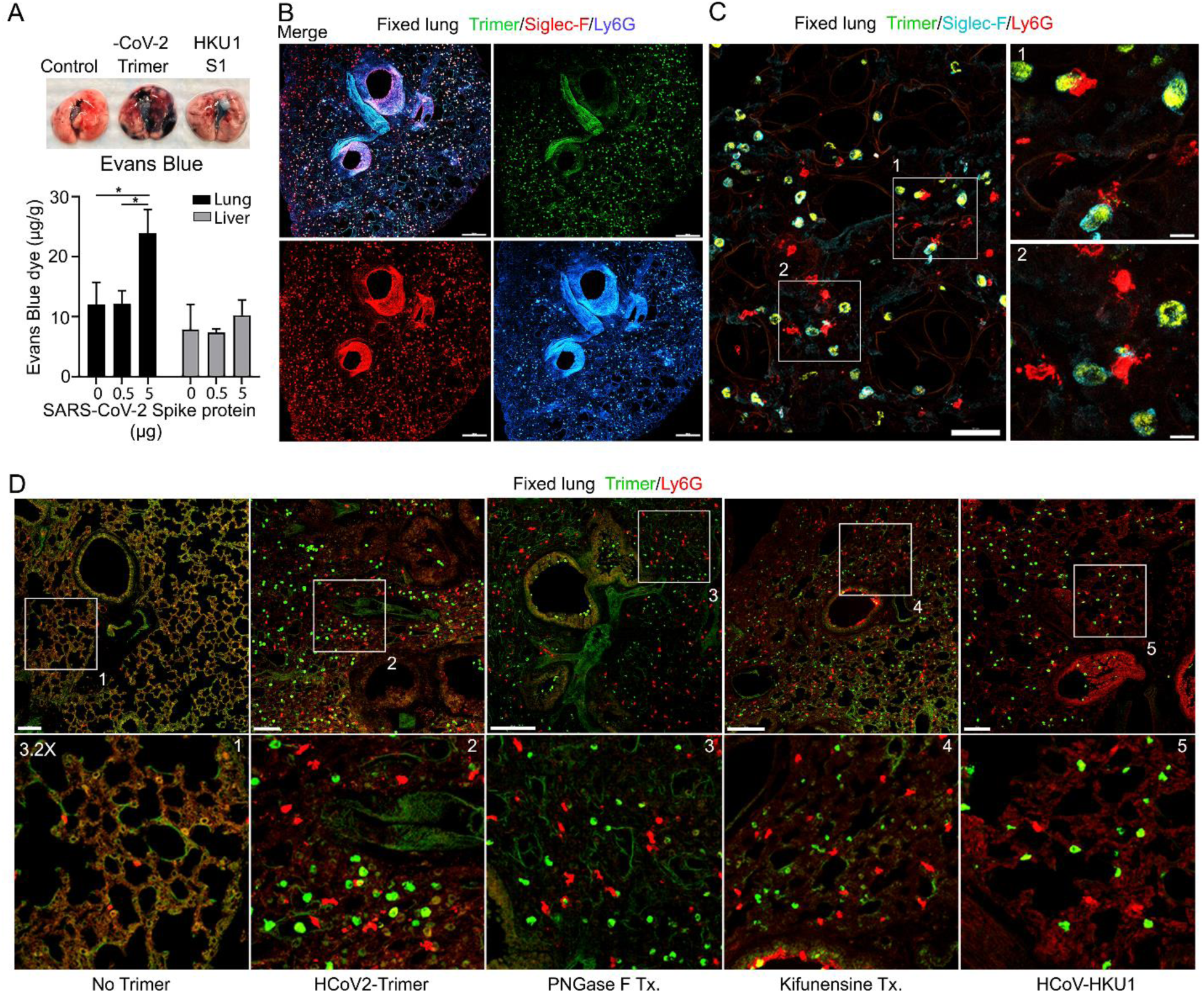
Increased lung vascular permeability and neutrophil localization following intranasal SARS-CoV-2 Spike protein. **(A)** Tissue photographs of lungs collected 2.5 hr post Spike protein (-CoV-2 Trimer), the S1 subunit of the human coronavirus (HKU1 S1), or saline instillation. Evans blue (200 ul of 5 mg/ml, PBS) was injected i.v. 1.5 hr post intranasal Spike administration. Lungs collected 1 hr after Evans blue injection. Evans blue amounts in lung and liver tissue are shown. **(B)** Confocal micrographs of lung collected at 18 hr post Spike protein. AMs and neutrophils detected with Siglec-F and Ly6G antibodies, respectively. Scale bar, 500 μm. **(C)** Confocal micrographs of lung collected at 3 hr post SARS-CoV-2 Spike protein instillation show SARS-CoV-2 Spike protein (green), neutrophils (red), and AMs (cyan). ROI-1 and −2 (boxes), show contact between AMs and neutrophils and are enlarged in the right panels. Scale bars, 50 and 10 μm. **(D)** Confocal micrographs of lung collected at 3 hr post each SARS-CoV-2 Spike proteins instillation show SARS-CoV-2 Spike protein (green) and neutrophils (red). From left to right of panels show saline (No Trimer), SARS-CoV-2 Spike protein (HCoV-2-Trimer), PNGase F treated SARS-CoV-2 Spike protein (PNGase F Tx.), SARS-CoV-2 Spike protein purified from Kifunensine treated cells (Kifunensine Tx.), and the S1 subunit of the human coronavirus HKU1 (HCoV-HKU1) Spike Protein. ROIs in each upper panel (boxes) are enlarged (3.2× magnification) in lower panels. Scale bars, 200 μm. *p < 0.05.

Next, we investigated the potential impact of modifying the glycans displayed by the spike proteins on lung vasculature homeostasis and neutrophil recruitment. Imaging fixed lung tissue from mice instilled with saline, SARS-CoV-2 Spike protein, glycan-deficient protein, high mannose protein, and the HCoV-HKU1 S1 subunit recombinant protein revealed enhanced neutrophil recruitment with each SARS-CoV-2 Spike protein preparation. We analyzed the neutrophil count in a 200 × 200 μm² area at six different locations. The surveyed areas of the SARS-CoV-2 Spike protein instilled mice had approximately a 6.5-fold higher neutrophil density compared to the saline-treated control mice. The high mannose version of the Spike protein recruited fewer neutrophils at the three-hour time point while the PNGase F treated version did not differ from the untreated trimer **(Figure 3D**, **Figure 3-figure supplement 1C)**.

### Neutrophil recruitment and damage in the cremaster muscle following local protein injection, and in the liver following intravenous injection

An exteriorized cremaster muscle is commonly used to intravitally image mouse neutrophil transmigration and interstitial motility (31). Although this model lacks Siglec F-positive macrophages, it is worth monitoring the effect of the SARS-CoV-2 Spike protein on neutrophils recruited into an inflammatory site. Within 3 hours local IL-1β injection recruits bone marrow neutrophils and leads to their transmigration into the nearby tissue. Co-injecting fluorescently labeled Ly6G and Gp1bβ antibodies identified neutrophils and platelets, respectively. Local PBS injection and intermittent imaging over a 6-hour time frame revealed occasional blood neutrophil and numerous flowing platelets **(Figure 4A, Video 1)**. Local intrascrotal injection of unlabeled SARS-CoV-2 Spike protein caused a modest recruitment of neutrophils at 4-6 hours post injection **(Figure 4A**, **Figure 4-figure supplement 1, Video 1)**. To better assess the long-term effect of the SARS-CoV-2 Spike protein, we co-injected Il-1β and waited until the following day to image. Typically, at 24 hours post Il-1β the inflammatory response is resolving, and the interstitial neutrophil numbers are declining from their peak **(Video 2)**. In contrast, each of the Spike protein preparations adversely affected neutrophil motility and morphology. Numerous neutrophils and neutrophil fragments localized along the blood vessel walls and were scattered within the interstitium **(Figure 4B, Video 2)**. We did not note any significant difference between the different SARS-CoV-2 Spike protein preparations as each caused neutrophil fragmentation and a decline in mobile neutrophils **(Figure 4C)**.

**Figure 4.**
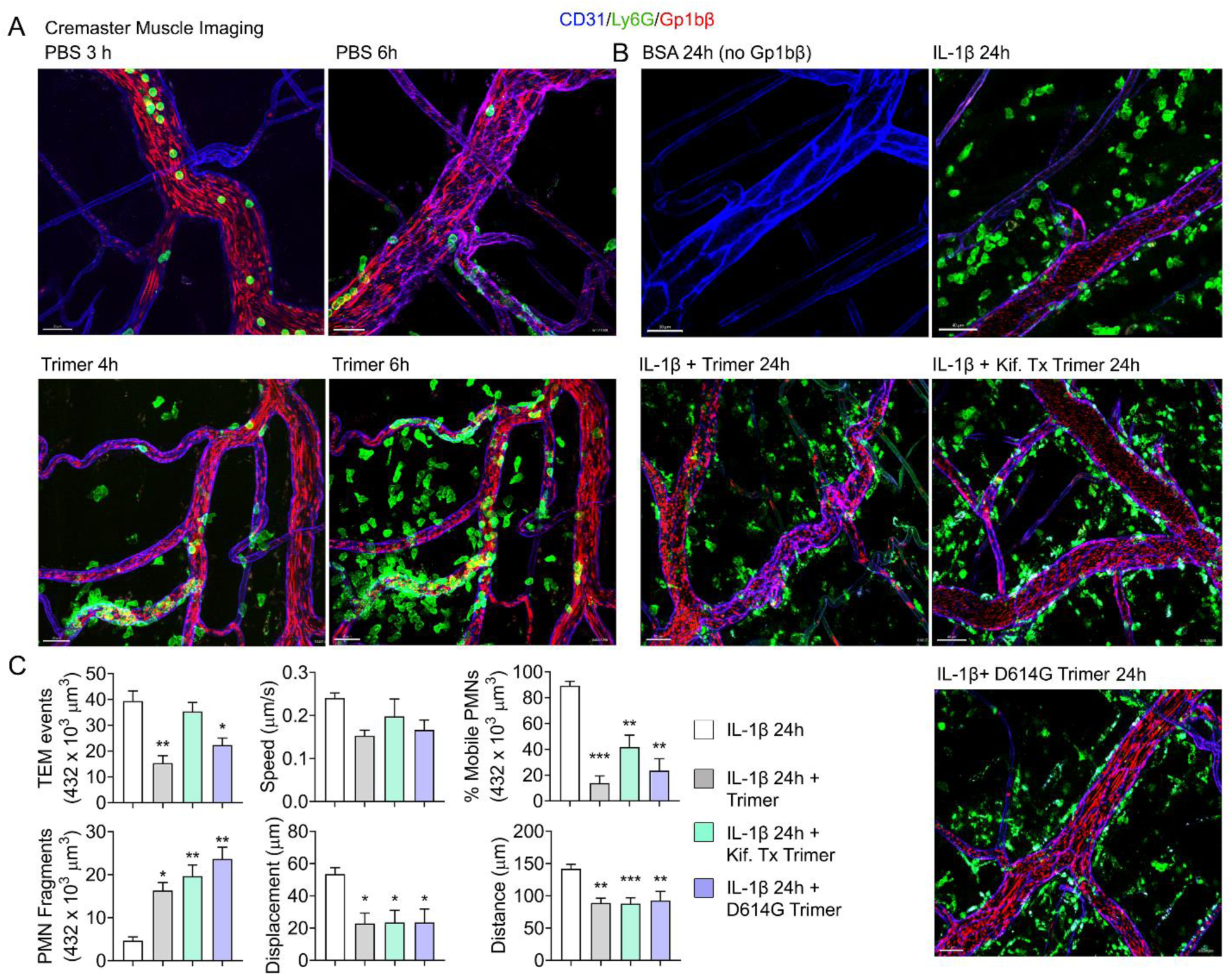
Mouse neutrophil damage following intrascrotal injections of Spike proteins. **(A & B)** Confocal snapshots of time-lapse movie of blood vessels in the cremaster muscle following PBS, BSA, Spike protein preparations (Trimer), IL-1β, or IL-1β plus Spike protein (5 µg) at indicated time points. Fluorescently labelled CD31, Ly6G, and Gp1bβ outlined the blood vessel endothelium, neutrophils, and platelets, respectively. Scale bars, 30 µm, except far right top and middle panels, 40 µm **(C)** Analysis of imaging data. The number of transendothelial migration (TEM) events recorded, neutrophil (PMN) fragments, and % mobile neutrophils in a defined imaging space over 20 min. Results of neutrophil tracking in the same defined volume including speed, displacement, and distance. Imaris software used for the tracking. *p < 0.05; **p < 0.005.

We also assessed the impact of injecting the labeled SARS-CoV-2 protein into the blood on the liver sinusoids using intravital microscopy. Snapshots at three hours post infusion revealed Spike protein outlined liver sinusoid endothelial cells co-localized with the CD31 delineated sinusoid endothelial membranes. Kupffer cells identified by F4-80 immunostaining rapidly acquired large amounts of the infused Spike protein **(Figure 4-figure supplement 2A, Video 3)**. The sinusoids, normally devoid of neutrophils, contained many Ly6G^+^ neutrophils. However, by 18 hours post infusion the amount of Spike protein outlining the sinusoids had declined as had the Kupffer cell associated material **(Figure 4-figure supplement 2B)**. The neutrophil infiltration had largely resolved suggesting that the inflammatory signals had declined. The origin of the white dots interspersed between the sinusoids is unknown, but they were not present at 3 hours post infusion despite identical imaging conditions. One possibility is that altered vascular permeability had allowed some labeled antibodies to leak into the liver parenchyma. To determine whether other endothelial beds also acquired intravenous administered Spike protein we examined the spleen, heart ventricle, and Peyer’s patches. At 3 hours post infusion endothelial cells at these three sites had acquired the labelled Spike proteins, most prominently in Peyer’s patches **(Figure 4-figure supplement 3A, B, C)**. Finally, we intravitally imaged the inguinal lymph node at 3 and 18-hour after local injection. Typically, locally injected material rapidly enters nearby afferent lymphatics for delivery to the lymph node where subcapsular sinus macrophages first encounter it (32). If it bypasses these macrophages lymph borne material flows into the lymph node medullary region, where medullary macrophages reside **(Figure 4-figure supplement 3D)**. At both the 3-hour time point (data not shown) and at 18 hours subcapsular macrophages showed little interest in the Spike protein, but medullary macrophages avidly acquired it. In contrast to the liver, we did not detect neutrophil recruitment into the lymph node at either time point after tail-base injection.

### Murine and human neutrophils NETosis following exposure to the SARS-CoV-2 Spike protein

Live cell imaging of lung sections following intranasal instillation of the Spike protein revealed ongoing neutrophil damage and likely NETosis (**Figure 5A**). Time-lapse images show several disrupted neutrophils near a Siglec-F positive AM along with Spike protein bound to a dying neutrophil. This observation along with the cremaster muscle imaging suggested direct neutrophil toxicity. Furthermore, a previous study had shown Spike protein induced neutrophil NETosis (33). To confirm that the Spike protein can trigger neutrophil NETosis and to compare different Spike protein preparations, we briefly exposed murine bone marrow neutrophils and assessed the % of cells undergoing cell death using a flow-based assay. Exposure to fMLP served as a positive control. The lowest concentration of Spike protein (0.1 µg) tested increased the number of dying neutrophils **(Figure 5B)**. Switching to human neutrophils purified from human peripheral blood, exposure to TNFα, LPS or PMA for 4 hours or overnight reduced their viability as expected. Addition of SARS-CoV-2 or the D614G mutant protein (1 µg/ml) decreased their viability at 4 hours and more dramatically upon overnight exposure. The high mannose versions of the Spike protein and D614G Spike had a slightly greater toxicity at 4 hours but had a similar impact in the overnight assay **(Figure 5C)**. Finally, we imaged human neutrophils overlaid on SARS-CoV-2 Spike protein treated A549 cells, a human lung epithelial cell line. Numerous dying neutrophils could be observed likely undergoing NETosis **(Figure 5D, Video 4)**. These results confirm that the SARS-CoV-2 Spike protein can cause neutrophil damage potentially exacerbating the inflammatory response.

**Figure 5.**
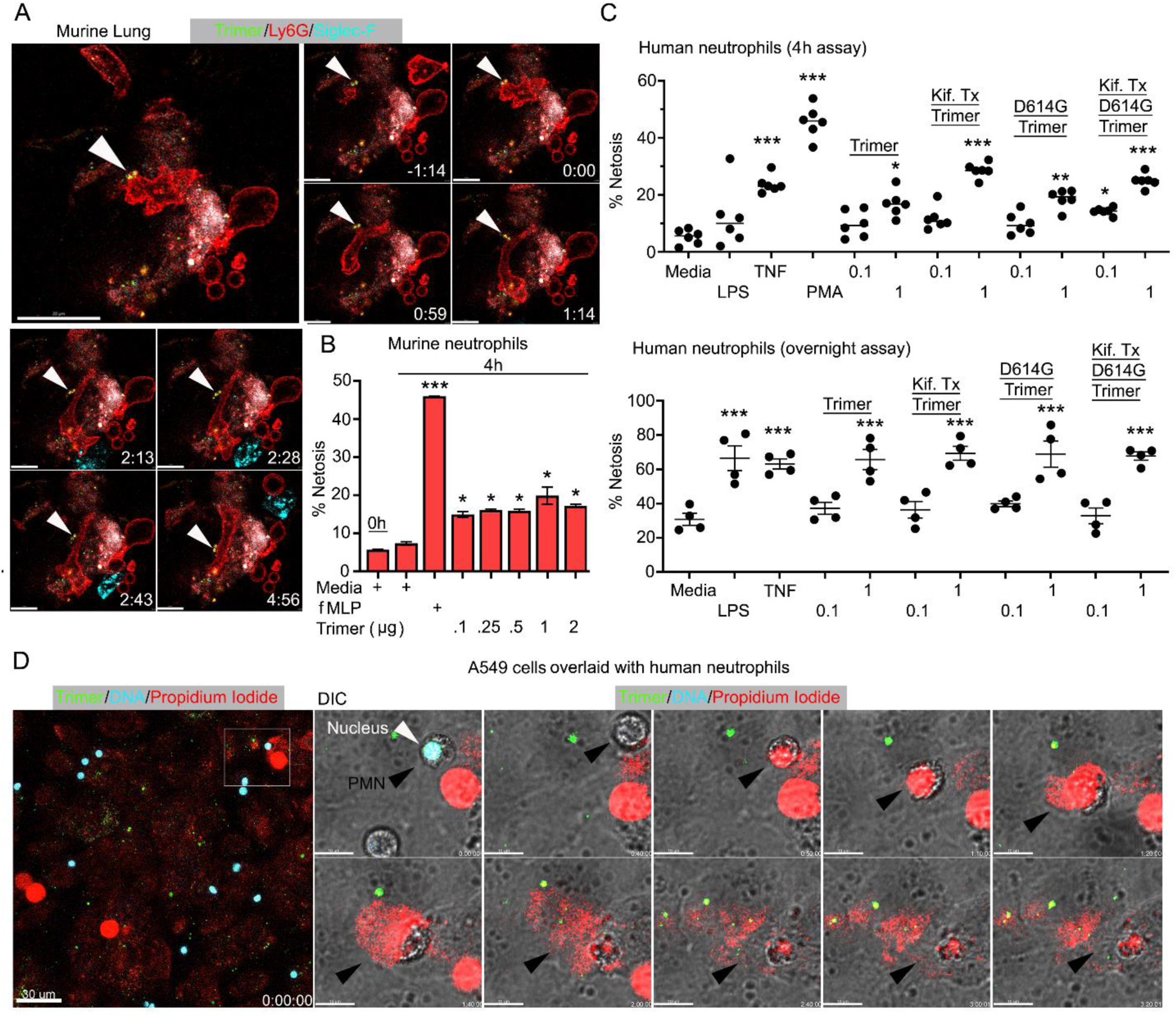
Neutrophil injury following exposure to SARS-CoV-2 Spike proteins. **(A)** Time-lapse images of a freshly sliced lung section 3 hr after SARS-CoV-2 Spike protein instillation. Neutrophils (Ly6G) and AMs (Siglec-F) were visualized by injected fluorescently tagged antibodies. Serial images show neutrophil behavior on Spike protein (Trimer) bearing cells. Arrowheads indicate Spike protein deposition. The time banner is set to 0:00 as neutrophil contacts Spike protein. Scale bars, 100, 25, and 20 μm. **(B)** Purified murine bone marrow neutrophils were exposed to increasing concentrations of Spike protein or fMLP (N-formyl-methionyl-leucyl-phenylalanine) for 4 hours or not. The percentage of neutrophils undergoing NETosis was measured by flow cytometry. **(C)** Purified human peripheral blood neutrophils were exposed to various Spike protein preparations, lipopolysaccharide (LPS), tumor necrosis factor-α (TNF) or phorbol myristate acetate (PMA). Graphs show the percentage of DAPI^+^Helix NP NIR^+^ cells after either 4 hr or overnight culture. Each point represents a different donor. **(D)** A still image of *in vitro* time-lapse movie shows exposed neutrophil DNA (red, propidium iodide), neutrophil nucleus (cyan, Hoechst), and SARS-CoV-2 Spike protein (trimer, green) bearing A549 cells. Plated A549 cells were treated with Spike protein (0.5 μg/ml) 24 hr prior to Hoechst-stained human neutrophil seeding. Propidium iodide (1 μg/ml) was added to the culture media and time-lapse images acquired every 10 min for 5 hr. Sequential DIC images overlay with fluorescent images show neutrophil NETosis. White arrowhead indicates a neutrophil’s intact nucleus (cyan, Hoechst). Black arrowheads delineate a neutrophil undergoing NETosis as detected by exposed DNA (red). Scale bars, 30 and 10 μm. *p < 0.05; **p < 0.01; ***p < 0.005.

### Human peripheral blood monocytes, B cells, neutrophils, and dendritic cells bind the SARS-CoV-2 Spike protein

Next, we assessed human peripheral blood leukocyte binding using the fluorescently labeled Spike protein. Binding assays were performed on ice to avoid endocytosis using Hanks balanced salt solution with added Ca^2+^ and Mn^2+^, or with EDTA to assess cation dependence. Representative CD4 T cell, B cell, Monocyte, or neutrophil flow patterns using either the Spike protein or the high mannose version are shown **(Figure 6A)**. The flow cytometry results demonstrated that cations enhanced leukocyte binding particularly so with the high mannose protein. The SARS-CoV-2 S1 protein bound better than did the HKU1 S1 protein, most evident with NK cells, neutrophils, monocytes, and dendritic cells **(Figure 6B)**. The SARS-CoV-2 Spike protein bound better than did the S1 protein with B cells, neutrophils, and monocytes exhibiting the best binding. The high mannose protein bound less well to the different leukocyte subsets. Surprisingly CD4 T cells poorly bound each of the proteins, irrespective of the presence or absence of cations. The D614G Spike protein bound better to neutrophils and monocytes than did the wild-type Spike protein (**Figure 6B**).

**Figure 6.**
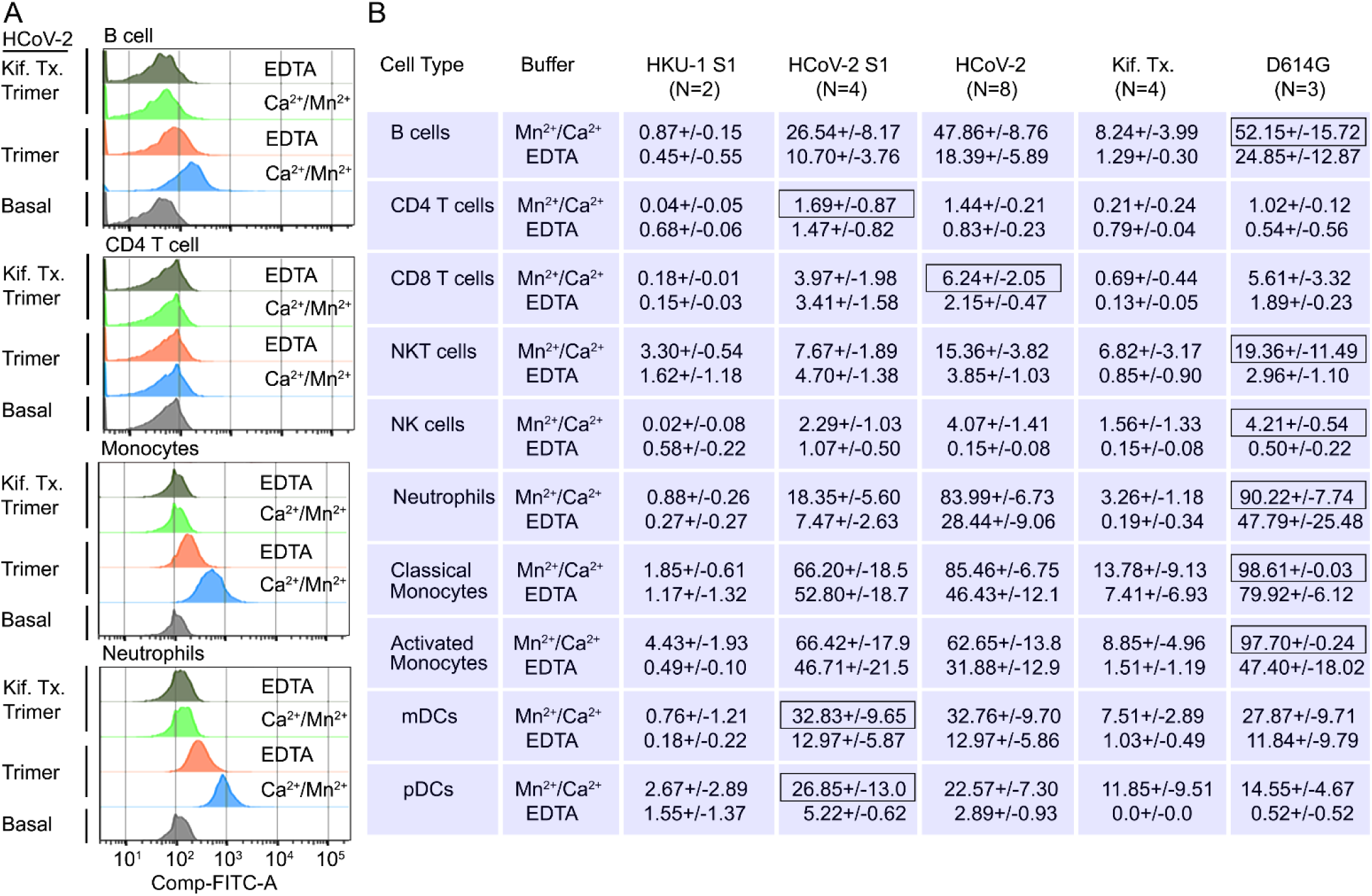
Binding of recombinant SARS-CoV Spike proteins to human peripheral blood leukocytes **(A)** Representative flow cytometry histograms of various cell populations prepared from whole blood incubated with labeled recombinant SARS-CoV-2 Spike protein (Trimer), or Spike protein from Kifunensine treated cell (Kif. Tx. Trimer), binding done in the presence of Ca^2+^/Mn^2+^ or EDTA. **(B)** Percentage of various cell populations prepared from whole blood that bound labelled HKU1 S1 protein (HKU-1 S1), SARS-CoV-2 S1 protein (HCoV-2 S1), SARS-CoV-2 Spike protein (HCoV-2 Trimer), Spike protein purified from Kifunesine treated cells (Kif. Tx. Trimer) or D614G Spike protein. Binding done in the presence of Ca^2+^/Mn^2+^ or EDTA and assessed by flow cytometry. Background fluorescent (unstained) subtracted from fluorescent signal. Data are from two-eight independent experiments. The recombinant protein that bound the highest % of cells of the different cell types is designated with black outline.

### Murine and human cells use Siglecs to help capture the SARS-CoV-2 Spike Protein

Since murine and human leukocytes lack significant ACE2 levels, the major entry receptor for SARS-CoV-1 and −2, they likely use other receptors to capture the Spike proteins. The strong co-localization with Siglec-F expressing murine AMs prompted an examination of the role of Siglec-F and other Siglecs in capturing the Spike proteins. Siglecs are transmembrane proteins that exhibit specificity for sialic acids attached to the terminal portions of cell-surface glycoproteins. Several viruses take advantage of sialic acid-Siglec interactions for cell targeting, spreading, and trans-infection (34–36). Initially, we established a bead assay to assess the binding to SARS-CoV-2 Spike proteins. We coupled the S1 domain protein, the SARS-CoV-2 stabilized Spike protein, or the PNGase F treated Spike protein and reacted the beads with fluorescently labeled antibody or different recombinant proteins. Flow cytometry analysis revealed strong binding of the Spike protein antibody, human ACE2, but not murine ACE2 as expected **(Figure 7A)**. Siglec-F also bound well, while the human Siglec-5 and Siglec-8 bound poorly despite being the structural and functional equivalents of Siglec-F, respectively (37). Of note the PNGase F treated fraction we used likely retained some N-linked glycans as it remained able to bind ACE2 although it lost Siglec-F binding (Figure 7A).

**Figure 7.**
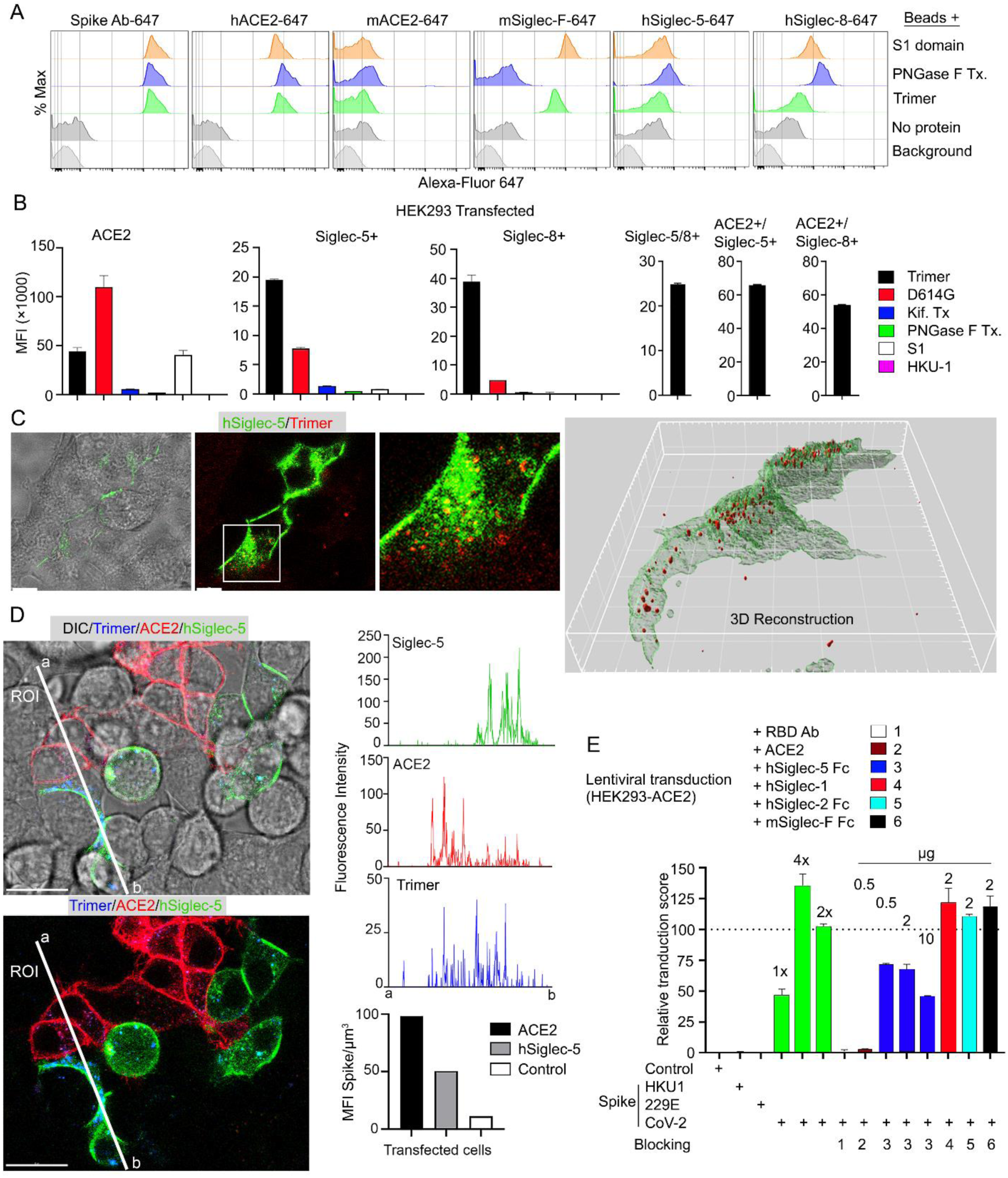
Role of Siglecs in SARS-CoV-2 Spike capture. **(A)** Histograms show mean fluorescence of conjugated recombinant proteins and antibody on nanobeads that are coated with indicated SARS-CoV-2 Spike proteins. Background signal (weak gray) measured with uncoated beads. No protein (gray) means that uncoated beads were incubated with indicated labelled recombinant protein or antibody. **(B)** Flow cytometry assessment of indicated Spike proteins binding to HEK293 cell permanently transfected with ACE2, various Siglecs, or a combination. Dual transfected HEK293 cells noted as Siglec-5/8+, ACE2/Siglec-5+, and ACE2/Siglec-8+. Labelled recombinant proteins were incubated with 5 × 10^3^ HEK293 cells on ice for 30 min. Data shown as mean fluorescent intensity. **(C)** DIC and confocal micrographs showing SARS-CoV-2 Spike (Trimer, red) protein acquisition by human Siglec-5-GFP transfected HEK293. ROI-1 in the middle panel is enlarged at the right panel. A 3D-reconstituted volume image of Siglec-5 transfected cell (far right), shows Siglec-5 mediated Spike protein acquisition. Scale bars, 30 and 10 μm. **(D)** DIC and confocal images show SARSCoV-2 Spike protein acquisition by human Siglec-5-GFP or ACE2-OFP transfected HEK293. Equal numbers of stable-transfectants expressing Siglec-5-GFP or ACE2-OFP, and non-transfectant were seeded together. Spike protein (Trimer, blue) was added 1 hr prior to imaging. In a region of interest (ROI) line ab fluorescence intensity of Siglec-5-GFP, ACE2-OFP and SARS-CoV-2 Spike trimer were analyzed. Fluorescent intensity graphs show fluorescence intensity of each signal in ROI. Amount of 3Dreconstructed volume of SARS-CoV-2 Spike protein in each stable cell and HEK293 cell were analyzed and plotted in graph. Graphs shows quantity of mean fluorescence intensity of SARS-CoV-2 Spike protein in unit volume (µm^3^) of indicated cell types. Scale bars, 20 μm. **(E)** Transduction of Spike protein expressing lentiviruses into ACE2/HEK293 transfectants. Lentiviruses envelop proteins: control (none), human coronavirus HKU1 Spike protein (HKU1), human coronavirus 229E Spike protein (229E), and SARS-CoV-2 Spike protein (CoV-2) are indicated. The volume of virus concentrates used for transduction is denoted as 1×, 2×, and 4× on the graph (bright green bars). Transduction scores were normalized to 2× transduction efficiency. Various recombinant proteins or antibody used for inhibition assay are indicated. Recombinant proteins or RBD neutralizing antibody were added 0.5 hr before lentivirus transduction. 3 days later GFP expression was quantitated by flow cytometry. Results are from 3 separate experiments. Statistics, 2x vs 2x + hSiglec-5 Fc (10 µg), p < 0.0005.

Despite the poor binding to the recombinant human Siglecs in the bead assay, we elected to test them in context of a human cell by expressing them in HEK293 cells. We chose HEK293 cells as they exhibited little Spike protein binding. We established ACE2, Siglec5, Siglec-8, Siglec-5/8, ACE2/Siglec-5, and ACE2/Siglec-8 expressing cell lines **(Figure 7B)**. The ACE2 transfected cells behaved as anticipated binding the original Spike protein and even better the D614G version; and binding the S1 protein. The high mannose and glycan deficient Spike proteins bound poorly. In contrast to the bead assay both the Siglec-5 and the Siglec-8 expressing HEK293 cells bound the Spike protein, although less efficiently than did the ACE2 expressing cells. They bound the D614G version less well, and very weakly bound the S1 domain protein. The high mannose and glycan deficient Spike proteins exhibited little binding to the Siglec expressing cells. Co-expression of Siglec-5 and −8 did not improve binding, although the ACE2/Siglec co-expressing cells bound the SARS-CoV-2 Spike protein slightly better than did ACE2 only cells. We also imaged the Siglec-5 expressing HEK293 with labeled protein **(Figure 7C)**. The imaging revealed efficient uptake and Spike protein endocytosis. Co-cultured ACE2 and Siglec-5 expressing cells both captured the Spike protein while HEK293 cells not expressing either protein failed to bind or uptake it **(Figure 7D, Video 5)**. To determine whether Siglec-5 or Siglec-8 could contribute to viral transduction, we produced lentiviral particles expressing the Spike protein from SARS-CoV-2, HKU1, or 229E and attempted to transduce the HEK293 ACE2 expressing cells. Successful transduction resulted in GFP expression, which we quantitated by flow cytometry and only occurred with the lentiviral particles with SARS-CoV-2 Spike protein incorporation **(Figure 7E)**. Both a receptor binding domain antibody and recombinant ACE2 blocked transduction. Recombinant human Siglec-5 partially inhibited although it required relatively high concentrations, while human Siglec-1, human Siglec-2, and murine Siglec-F had no impact on the transduction frequency.

### Increased TNF-**α** and IL-6 production by human macrophages exposed to the SARS-CoV-2 Spike proteins

Dysregulated cytokine production contributes to the pathogenesis of severe COVID-19 infections (38, 39). To assess whether the Spike proteins can affect macrophage cytokine profiles we cultured human monocytes using conditions that generate M0, M1, or M2 macrophages and treated them with different Spike protein preparations. We collected cell supernatants two days later and measured a panel of known macrophage derived cytokines by ELISA **(Figure 8)**. We found little induction of IL-1β in the cell supernatant indicating that the Spike proteins alone did not trigger the inflammasome activation in these macrophages. In contrast, we did observe significant increases in TNF-α and IL-6 secretion by the original SARS-CoV-2 Spike protein treated macrophages irrespective of whether it was produced in CHO or HEK293F cells. The spike protein produced in HEK293F is predicted to contain more complex carbohydrates than that expressed in CHO cells. While both preparations increased IL-6 and TNF-α production, the spike protein produced by HEK293F cell elicited more TNF-α and less IL-6 compared with the CHO produced spike protein. The spike protein produced in CHO cells treated with Kifunesine elicited a similar cytokine profile as did the spike protein produced in CHO cells in the absence of Kifunesine. The addition of the HKU1 S1 protein did not modify cytokine production by any of the macrophage subsets.

**Figure 8.**
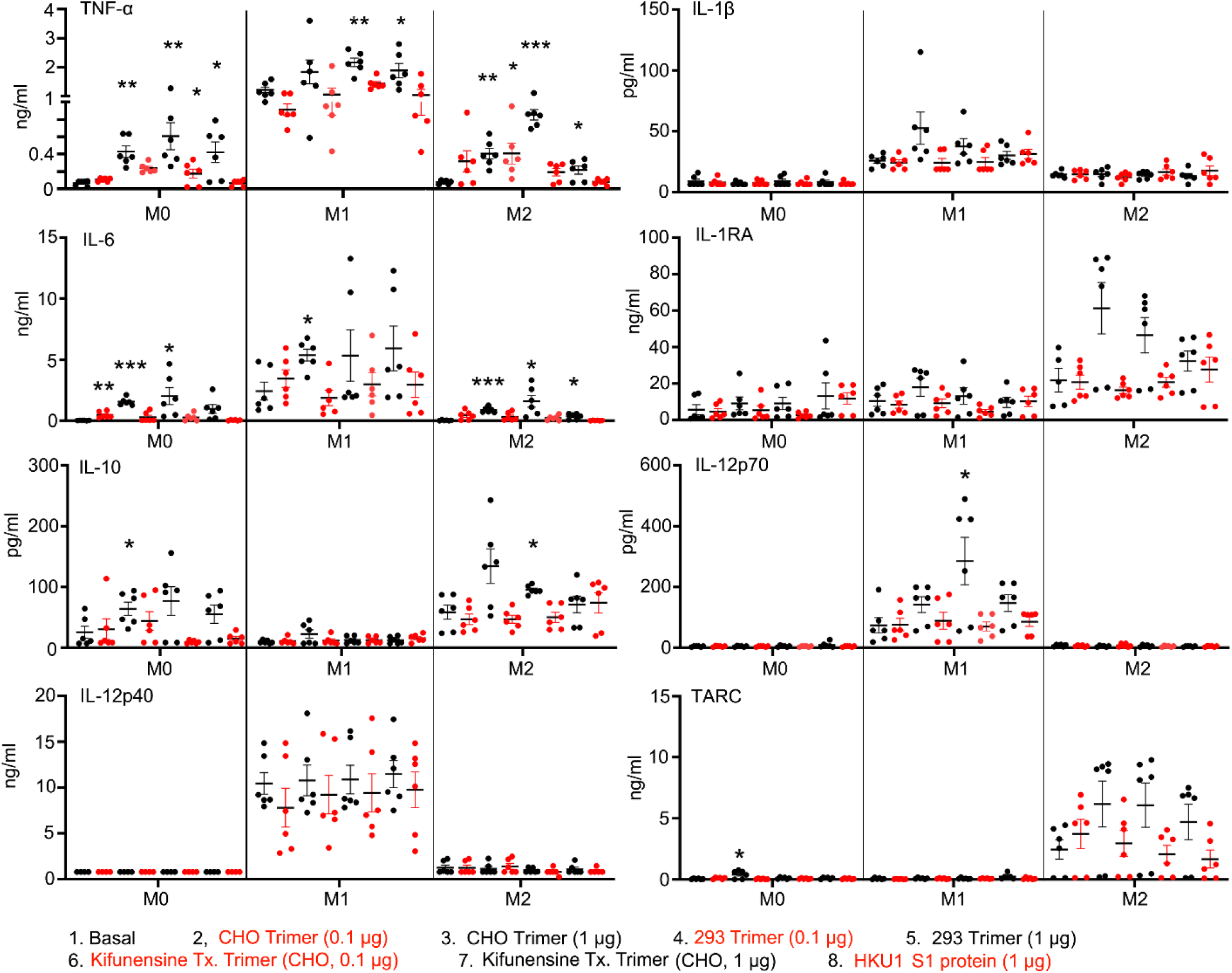
Macrophage cytokine profiles following exposure to different Spike proteins. Quantification of macrophage-associated cytokines in response to indicated Spike proteins. The graph shows the TNF-α, IL1β, IL-6, IL-12p70, IL-12p40, and TARC quantification of M0, M1, or M2 Macrophage cultures following stimulation, or not, with SARS-CoV-2 Spike proteins derived from CHO cells (CHO Trimer; 0.1µg/ml or 1 µg/ml), or 293F cells (293 Trimer; 0.1 µg/ml or 1 µg/ml), or purified Spike protein from Kifunensine treated CHO cells (Kifunensine Tx. Trimer; 0.1 µg/ml or 1 µg/ml), or Human coronavirus HKU1 S1 protein (HKU1 S1 protein; 1 µg/ml) for 48 hr. The error bars denote the mean ± SEM. Two-way ANOVA was used to compare treated samples to basal M0, M1, or M2 population (n = 6, *p < 0.05; **p < 0.01; ***p < 0.001).

## Discussion

This study analyzed the cell tropism of the SARS-CoV-2 Spike protein and some of its variants using fluorescently labeled proteins, flow cytometry, and mice to assess *in vivo* binding. The intranasal administration of SARS-CoV-2 Spike protein to mice led to its rapid uptake by Siglec-F positive AMs and an increased the number of neutrophils, monocytes, and dendritic cells in the lung 18 hours after instillation. Modifying the carbohydrate content of the Spike protein or using a Spike protein with a D614G mutation slightly altered the cell recruitment pattern. The high mannose Spike protein purified from the Kifunensine treated CHO cells increased the number of lung macrophages at 18 hours despite a lower % of macrophages having retained it. As alveolar macrophages express the mannose receptor (MR, CD206), a member of the C-type lectin (CLEC) family, perhaps the high mannose Spike protein is more rapidly endocytosed and degraded accounting for a lower percentage of cells retaining it. This receptor binds high-mannose structures present on the surface of pathogens helping to neutralize them by phagocytic engulfment. Other cell types including immature dendritic cells (DCs), and endothelial cells in hepatic, splenic, lymphatic, and dermal microvasculature express the mannose receptor and would be expected to be targeted by the high mannose Spike protein. We confirmed Siglec-F as a capturing receptor, as it efficiently bound to the S1 domain, and the SARS-CoV-2 Spike protein. Yet alveolar macrophages employ additional receptors to capture the Spike protein as they still accumulated the PNGase F treated Spike protein, which had lost binding to Siglec-F.

Arguing against a role for mSiglec-1, CD169^+^ lung macrophages and CD169^+^ subcapsular sinus macrophages failed to accumulate the Spike protein *in vivo*. The closest human paralog of mouse Siglec-F is hSiglec-8 (40). While expressed on human eosinophils and mast cells, human AMs apparently lack it. In contrast, human AMs do express Siglec-5 (37). Along with its paired receptor, hSiglec-14, Siglec-5 can modulate innate immune responses (41). When tested in a bead binding assay, in contrast to Siglec-F, neither hSiglec-5 or −8 bound the recombinant Spike protein, yet their expression in a cellular context allowed binding. Additionally, recombinant hSiglec-5 partially inhibited the transduction of Spike protein bearing viral like particles into ACE2 expressing HEK293 cells. Evidently the recombinant protein binding assay we employed did not fully capture the properties of hSiglec-5 or hSiglec-8 in a cellular context. A recent study of human lungs revealed scant alveolar ACE2 expression, highlighting the importance of alternative Spike protein receptors (42). *Ex vivo* infected human lungs and COVID-19 autopsy samples showed AMs positive for SARS-CoV-2 and single-cell transcriptomics revealed nonproductive virus uptake, and activation of inflammatory and anti-viral pathways. Another study which analyzed pulmonary cells from COVID-19 patients found that myeloid cell C-type lectin engagement induced a robust proinflammatory responses that correlated with COVID-19 severity (43). The robust Spike binding to AMs and their potential roles in COVID-19 lung infection warrants a further exploration of their Spike protein binding partners.

The intranasal administration of the Spike protein resulted in an increase in lung vascular permeability and local tissue damage. While the mechanisms are unknown, infectious agents that target AMs can trigger the secretion of factors that disrupt the integrity of the pulmonary microvascular cell barrier. For example, Porcine reproductive and respiratory syndrome virus infected AMs trigger transcriptional changes in genes in co-cultured microvascular endothelial cells that reduce vascular integrity (44). Lung endothelial cells may also be directly targeted (45). Spike protein/ACE2 interactions reduce endothelial cell ACE2 expression, which can alter vascular permeability. Yet, the low affinity of mouse ACE2 for the SARS-CoV-2 Spike protein presumably precludes this mechanism in this study. Also arguing against a role for ACE2 in our model, the responses to intranasal Spike protein in human ACE2 transgenic mice resembled those of wild type mice. The Spike protein RGD (Arginine-Glycine-Aspartic Acid) motif can also bind α_V_β_3_ integrins present on endothelial cells (46). This affects VE-cadherin function and vascular integrity. However, we did not detect any Spike protein on the pulmonary blood vessels following intranasal administration although a low level of binding may have escaped visualization. In sum, the rapid and intense uptake of Spike protein by AMs supports a role for these cells in the localized neutrophil recruitment, nearby tissue damage, and increased vascular permeability that follows its intranasal administration.

The intravenous administration of the Spike protein, led to its accumulation by Kupffer cells in the mouse liver. This uptake was accompanied by a transitory increase in liver sinusoid neutrophils. Human Kupffer cells express LSECtin (liver Siglec and lymph node sinusoidal endothelial cell C-type lectin, CLEC4G) a C-type lectin receptor encoded within the L-SIGN/DC-SIGN/CD23 gene cluster. LSECtin acts as a pathogen attachment factor for Ebola virus and the SARS coronaviruses suggesting that it contributed to the Kupffer cell uptake we noted (47). The liver sinusoid endothelial cells also avidly accumulated the Spike protein. These cells express another C-type lectin receptor L-SIGN (CD209L/CLEC4M), which has been shown to interact in a Ca^2+^-dependent manner with high-mannose–type N-glycans on the SARS-CoV-2 Spike protein (48). Based on our mouse studies, blood borne SARS-CoV-2 Spike proteins, viral particles, or exosomes bearing the Spike protein are likely to accumulate in the liver during severe or prolonged COVID-19 infection.

Local injection of the Spike protein in the region of the cremaster muscle recruited low numbers of neutrophils, but co-administration with IL-1β caused severe neutrophil damage. The Spike protein also proved toxic to purified mouse and human neutrophils cultured *ex vivo* in its presence. Most human neutrophils bound the recombinant Spike protein. The presence of cations in the binding buffer substantially enhanced the binding suggesting that a cation dependent receptor accounted for a significant portion of the binding. While the high mannose version bound neutrophils less well, in the NETosis assay it produced a similar level of neutrophil cell death. Neutrophils likely employ several different receptors to capture the Spike protein. Human neutrophils express several C-type lectin receptors including CLEC5A, which has been implicated in SARS-CoV-2 triggered neutrophil NETosis (49). They also express Siglec-5 (37), which bound the Spike protein when expressed in HEK293 cells.

We also assessed the binding of various Spike protein preparations to human peripheral blood mononuclear cells and their impact on cytokine production by M0, M1, and M2 human monocyte derived macrophages. Monocytes, neutrophils, and B cells bound the full-length trimer best, while T and NK cells exhibited little binding. All the cellar subsets bound more of the SARS-CoV-2 Spike protein in the presence of cations arguing for a major role for cation dependent receptors. The high mannose Spike protein bound less well to all tested cell types and its binding exhibited the highest cation dependence. Among the eight cytokines measured in the supernatants conditioned by human monocyte derived macrophages treated with Spike proteins only TNF-α and IL-6 were elevated compared to untreated or HKU1 S1 treated cells. We found little or no change in IL-1β levels suggesting that the tested Spike protein preparations did not activate inflammasomes. Nor did we find any material difference between recombinant Spike protein prepared from HEK293F and CHO cells. Further studies are needed to identify the monocyte, neutrophil, and B cell receptors that account for the Spike protein binding and additional functional studies to assess the consequences of that binding.

Several conclusions can be drawn from this study. First, instilled SARS-CoV-2 Spike protein and VLPs bearing the Spike protein rapidly accumulated on mouse AMs suggesting a similar uptake by human alveolar macrophages during an active SARS-CoV-2 infection. In mice, the nasal instillation of the Spike protein is accompanied by lung leukocyte infiltration. Second, murine alveolar macrophages likely use Siglec-F as a capturing receptor although other receptors contribute. Third, following nasal instillation of the Spike protein, the integrity of the pulmonary vasculature weakens. Fourth, resident and recruited neutrophils suffer, either directly targeted by the Spike protein, or secondarily due to the inflammatory response. A similar scenario would be expected during SARS-CoV-2 infection, following direct instillation of the Spike protein in the upper airway of humans, or following an intranasal vaccine that directs the synthesis of the SARS-CoV-2 Spike protein or fragments. Fifth, blood monocytes, B cells, neutrophils, and dendritic cells efficiently bind the Spike protein, while only a low percentage of CD8 T cells and even fewer CD4 T cells do. Spike protein binding to human monocyte derived macrophages increases the TNF-α and IL-6 secretion. Sixth, altering the glycan composition of the Spike and the D614G mutation had surprisingly little impact on the initial inflammatory response. Seventh, viral particles and soluble Spike protein that spills into the blood during SARS-CoV-2 infection or following immunization will likely be cleared by liver Kupffer cells but will also target blood vessel endothelial cells in multiple organs. Finally, injection of the Spike protein as occurs following vaccination rapidly delivers it to the draining lymph node via afferent lymphatics. Surprisingly, lymph node medullary macrophages, which largely serve a degradative function, preferentially uptake the Spike protein. Designing a less toxic Spike protein immunogen that targets subcapsular sinus macrophages might better deliver the immunogen to B cells, thereby eliciting a more efficacious antibody response.

## Material and Methods

### Mice

C57BL/6 and K18-ACE2 transgenic mice were obtained from Jackson Laboratory. All mice were used in this study were 8-12 weeks of age. Mice were housed under specific-pathogen-free conditions. All the animal experiments and protocols used in the study were approved by the NIAID Animal Care and Use Committee (ACUC) at the National Institutes of Health.

### Cells

To isolate mouse lung cells, lungs were carefully collected and gently teased apart using forceps into RPMI 1640 media containing 2 mM L-glutamine, antibiotics (100 IU/ml penicillin, 100 μg/ml streptomycin), 1 mM sodium pyruvate, and 50 μM 2-mercaptoethanol, pH 7.2. The tissue was then digested with Liberase Blendzyme 2 (0.2 mg/ml, Roche Applied Science) and DNase I (20 μg/ml) for 1hr at 37°C. The proteases were inactivated by adding 10% fetal bovine serum and 2 mM EDTA and the cell disaggregated by passing them through a 40 μm nylon sieve (BD Bioscience). Single cells were then washed with 1% Bovine serum albumin (BSA) in PBS and blocked with anti-Fcγ receptor (BD Biosciences). Human peripheral blood mononuclear cells (PBMC) were purified from whole blood by density gradient centrifugation (FicollPaque^TM^, Miltenyi Biotec). Whole blood was collected from healthy donors through an NIH Department of Transfusion Medicine (DTM)-approved protocol (Institutional Review Board of the National Institute of Allergy and Infectious Diseases [NIAID]). Neutrophil and monocyte cell population were each obtained by negative selection (>97% purity, Stem Cell Technologies). To generate human monocyte-derived macrophages, purified monocytes were treated for 7 days with 50 ng/ml human recombinant M-CSF (PeproTech) in RPMI medium supplemented with 10% heat inactivated fetal bovine serum, and 1mM Sodium Pyruvate, 100U Penicillin-Streptomycin (Gibco, ThermoFisher Scientific) in ultra-low attachment culture 100mm dishes. On day 7 the mature macrophages were collected and verified to be more than 90% CD68^+^ by flow cytometry.

### Reagents

See reagent table.

### Flow Cytometry

Single cells were re-suspended in 0.1% fatty acid free BSA-Hanks’ Balanced Salt Solution (HBSS) with 100 μM CaCl2 and 1 mM MnCl2 unless otherwise specified. Buffer without divalent cations included 10 mM EDTA. The cells were stained with fluorochrome-conjugated antibodies against various cell surface markers or with different fluorochrome-conjugated proteins, which are listed in the resource and reagent tables in the supplement. LIVE/DEAD™ Fixable Aqua Dead Cell Stain Kit, LIVE/DEAD™ Fixable NearIR Dead Cell Stain Kit or LIVE/DEAD™ Fixable Yellow Dead Cell Stain Kit (Thermo Fisher) were used in all experiments to exclude dead cells. Compensation was performed using AbC™ Total Antibody Compensation Bead Kit (Thermo Fisher) and ArCTM Amine Reactive Compensation Bead (Thermo Fisher) individually stained with each fluorochrome. Compensation matrices were calculated with FACSdiva software. Data acquisition included cell number count was done on a FACSCelesta SORP (BD) flow cytometer and analyzed with FlowJo 10.8.x software (Treestar).

### Human macrophage polarization and cytokine profiling

To polarize human macrophages, monocyte derived macrophages were plated in 48 well plates at 5 × 10^4^ cells per well in 500µl RPMI medium and allowed to rest 2hours at 37°C before treat of cytokines or LPS. The human M0 macrophages were either left untreated or treated for 48h to induce macrophage polarization: (i) for M1 polarization with 10 ng/mL LPS (E055:B55; Sigma-Aldrich) and 20 ng/mL IFNγ (PeproTech), (ii) for M2 polarization with 50 ng/ml human recombinant M-CSF, 20 ng/ml human recombinant IL-4, and 20 ng/ml human recombinant IL-13 (PeproTech). Human macrophages were then cultured alone or with 0.1ug, or 1ug SARS-CoV-2 Spike proteins for 48 hr. The cultured supernatants were collected, and cytokine levels determined using the bead-based immunoassays LEGENDplex™ Human Macrophage/ Microglia Panel (Biolegend). The assays were performed in 96-well plates following the manufacturer’s instructions. For measurements a FACSCelesta SORP flow cytometer (BD Biosciences) was employed, and data were evaluated with the LEGENDplex™ Data Analysis software.

### Production and purification of recombinant SARS-CoV-2 Spike proteins (see Figure 1-figure supplement 1)

SARS-CoV-2 Spike expression vectors were purchased from SinoBiological (see reagent table). Endotoxin-free plasmids were prepared by Alta Biotech and provided at a concentration is 5mg/mL. Transfections were carried out using CHO Freestyle cells (Invitrogen). 1.24mg of plasmid DNA was transfected into 90 million cells were using a MaxCyte Electroporation Transfection System. A detailed protocol is available at https://maxcyte.com/atx. Culture supernatants were harvested on day 6 and clarified by centrifugation, followed by filtration through a 0.45µm filter, followed by the addition of EDTA-free protease inhibitor cocktail tablets (Roche). Culture supernatants were then dialyzed overnight at 4 °C in HBS (150mM NaCl, 10mM HEPES, pH 8.0) using 10kDa MWCO Slide-A-Lyzer dialysis cassettes (Thermo Scientific). Supernatants were passed over a 10 ml StrepTrap HP column (Cytiva) at 1 ml/min using an ÄKTA pure 150 purifier (Cytiva), maintained at 4°C. Bound protein was eluted with elution buffer (2.5mM desthiobiotin, 100mM Tris-Cl, 150mM NaCl, 1mM EDTA, pH 8.0). Peak fractions were pooled and concentrated using 30kDa MW CO Amicon centrifugal concentrators (Millipore). Trace endotoxins were removed by two sequential Triton X114 extractions, followed by passage through an HiPPR detergent removal column (Thermo Fisher). Following the removal of endotoxin, the remaining contamination was assessed using the Pierce™ Chromogenic Endotoxin Quant Kit (Thermo Fisher), based on the amebocyte lysate assay. The endotoxin level in the purified recombinant protein preparation is below 1.0 EU/ml, which closely aligns with the levels specified by the company for recombinant proteins. Protein concentrations were determined by a BCA Protein assay (Thermo Fisher).

### Viral like particles and lentivirus preparation

SARS-CoV-2 Spike protein incorporated NL4.3-GFP VLPs were produced by transfecting HEK293T cells with full length SARS-CoV-2 Spike-S and HIV-1 NL4-3 Gag-iGFP ΔEnv (12455, NIH AIDS Reagent Program) at a ratio of 1:2.5 using a previously reported method (32). EGFP control lentiviruses (no Spike protein) were produced by transfecting HEK293T cells with pCMV-dR8.2 dvpr (packaging) and pLentipuro3 TO V5-GW EGFP-Firefly Luciferase (reporter). Coronavirus Spike proteins were incorporated by co-transfection with SARS-CoV-2 Spike-S, HCoV-229E Spike, or HCoV-HKU1 Spike expressing plasmids. Eighty percent confluent HEK293T in six well plates (2.5 ml/well) were transfected with 5 μg of plasmid diluted in Opti-MEM with a 1:4 (DNA/reagent) dilution of TransIT-293 Transfection Reagent. The media was harvested 64 hr later and centrifuged at 500 ×g for 10 min at 4 °C. The supernatants were collected and mixed with Lenti-X Concentrator (Takara Bio USA, Inc) at a 1:3 ratio. The mixture was placed at 4 °C overnight and centrifuged the following day at 1500 ×g for 45 min. Fluorescent VLPs were directly counted and measured by BD FACSCelesta™ flow cytometer. The forward scatter (FSC) detector (Photodiode with 488/10 BP filter) and side scatter (SSC) detector (photomultiplier tube (PMT) with 488/10 BP filter) were tuned to detect voltages up to 530 for FSC and 220 for SSC. The NL4.3-GFP VLPs were distinguished from noise and non-VLP particles by GFP signals. NL4.3-GFP VLP Spike protein incorporation was tested using human ACE2 expressing HEK293 cells (ACE2/HEK293). Ten thousand VLPs were incubated with 1×10^4^ ACE2/HEK293 cells in 1× HBSS (contains; 1 mM Ca^2+^, 2 mM Mg^2+^, 1 mM Mn^2+^ and 0.5% fatty acid free BSA) buffer at 4 °C for 30 min. Binding of the VLPs to ACE2/HEK293 cells was measured with BD FACSCelesta™. The 50% lentivirus transduction dose was determined using percentile (50%) of GFP positive ACE2/HEK293 cells 3 days after transduction. The pelleted VLPs or lentiviruses were suspended in PBS and frozen in single use aliquots.

### Lentivirus transduction assay

96-well plates were seeded with 200 µl of ACE2/HEK293 cells (1.5×10^4^ cells/ml). After a 24 h incubation, recombinant proteins or neutralizing antibody was added. Lentiviral particles capable of transducing 50% of the cells (CoV2-lenti, HKU1-lenti, and 229E-lenti) were added (∼25 µl) 30 min later. Following a three-day culture, the ACE2/HEK293 cells were harvested and GFP expression levels determined by BD FACSCelesta™ flow cytometry. Each group contained 2-5 replicates.

### Measurement of vascular permeability

Pulmonary vascular permeability was measured by i.v. administration of Evans blue dye (0.2 ml 0.5% in PBS) (50). One and half hours after Spike protein nasal administration, Evans blue dye was injected intravenously. After another 1.5 hours the mice were perfused with PBS, and lungs and livers were harvested, and dye extracted in formamide overnight at 55°C. Dye concentrations were quantified by measuring absorbance at 610 nm with subtraction of reference absorbance at 450 nm. The content of Evans blue dye was determined by generating a standard curve from dye dilutions.

### NETosis assays

Mouse bone marrow derived- or human neutrophils (> 97% purity) were resuspended in RPMI 1640 media (1×10^6^/ml) and incubated at 37°C in 5% CO_2_ for 30 min. To induce NETosis, the cells were exposed to N-formylmethionyl-leucyl-phenylalanine (fMLP, 1 µM); lipopolysaccharide (LPS, 10 ng/ml); tumor necrosis factor-α (TNFα, 20 ng/ml); phorbol myristate acetate (PMA, 30 nM), or four different SARS-CoV Spike protein preparations (0.1µg/ml or 1µg/ml) for 4 hours to overnight at 37℃. The cultures were terminated by adding 4% PFA for 15 min. Helix NP^TM^ NIR (0.1 μM; Biolegend) and DAPI (0.3 nM; Biolegend) were added to detect NETs. Data acquisition (Helix NP^TM^ NIR^+^ DAPI^+^ Cells) was done on FACSCelesta SORP (BD) flow cytometer and analyzed with FlowJo software (Tree star). *In vitro* imaging of NETosis was performed using a modified protocol from a previous report (51). The A549 cells were plated 48 hr before imaging at 60% cell confluent. Fluorescently labeled SARS-CoV-2 Spike protein (Alexa Fluor 488; 0.5 μg/ml) was added to the A549 cells 24 hr before human neutrophil seeding. Human neutrophils were purified from whole blood by EasySep™ Direct Human Neutrophil Isolation Kit (Stem cell). Purified human neutrophils were stained with Hoechst prior to seeding on SARS-CoV-2 Spike protein pre-treated A549 cells in Propidium Iodide (PI, 1 μg/ml) containing culture media. Time-lapse images were acquired with a Leica SP8 inverted 5 channels confocal microscope (Leica Microsystems) equipped with 40× oil objective, 0.95 NA (immersion medium used distilled water). The temperature of air (5% CO_2_) was maintained at 37.0± 0.5°C. Time interval between frames were set as 10 min for 5 hr acquisition. The loss of nucleus of human neutrophils was detected by the loss of Hoechst signal and extracellular DNA released by NETosis was labeled by PI signals.

### Fluorescent nanobead binding assay of SARS-CoV-2 Spike protein

FluoSpheres™ NeutrAvidin™-Labeled Microspheres, 0.2 µm, yellow-green fluorescent (505/515), 1% solids (Cat# F8774, Thermo Fisher Scientific Inc.) was used as a nanobead platform for SARS-CoV-2 Spike protein conjugation. Fluorescent nanobead were directly counted and measured by BD FACSCelesta™ flow cytometer. A flow cytometer equipped with forward scatter (FSC) detector (Photodiode with 488/10 BP filter) and side scatter (SSC) detector (photomultiplier tube (PMT) with 488/10 BP filter) was tuned to detect voltages up to 550 for FSC and 230 for SSC. Million counts of nanobeads were conjugated with 0.5 μg of SARS-CoV-2 Spike protein by interaction of Neutravidin on beads and Strep Tag II on recombinant proteins. Nanobeads and SARS-CoV-2 Spike proteins in PBS were coupled at room temperature for one hour and washed with 1 ml of PBS. Coupled nanobeads spun were down with benchtop centrifuge speed at 20,000 ×g, 4°C for 20 min. The nanobeads pellet was suspended with PBS concentration at 2 × 10^4^/μl. SARS-CoV-2 Spike protein couplings to nanobeads were tested with Alexa Fluor 647 conjugated SARSCoV-2 Spike neutralizing antibody (Cat# 40591-MM45, SinoBiological). To test binding ability of various recombinant proteins (human ACE2, human Siglec-5, human Siglec-8, mouse ACE2, and mouse Siglec-F) were directly conjugated with Alexa Fluor 647. SARS-CoV-2 Spike protein conjugated nanobeads (2× 10^4^ counts) in 20 μl of 1× HBSS (contains; 1 mM Ca^2+^, 2 mM Mg^2+^, 1 mM Mn^2+^ and 0.5% fatty acid free BSA) buffer were incubated with 0.2 μg of fluorescent recombinant proteins at room temperature for 30 min. Fluorescent antibody or recombinant protein biding on fluorescent nanobeads were directly counted and measured by BD FACSCelesta™ flow cytometer.

### Thick section immunohistochemistry and confocal microscopy

Immunohistochemistry was performed using a modified method of a previously published protocol (32, 52). Briefly, freshly isolated lungs were fixed in newly prepared 4% paraformaldehyde (Electron Microscopy Science) overnight at 4 °C on an agitation stage. Fixed lungs were embedded in 4% low melting agarose (Thermo Fisher Scientific) in PBS and sectioned with a vibratome (Leica VT-1000 S) at a 50 μm thickness. Thick sections were blocked in PBS containing 10% fetal calf serum, 1 mg/ml anti-Fcγ receptor (BD Biosciences), and 0.1% Triton X-100 (Sigma) for 30 min at room temperature. Sections were stained overnight at 4°C on an agitation stage with Ly6G, Siglec-F, CD169, CD31, and F4/80, and labelled WGA. Stained thick sections were microscopically analyzed using a Leica SP8 confocal microscope equipped with an HC PL APO CS2 40× (NA, 1.30) oil objective (Leica Microsystem, Inc.) and images were processed with Leica LAS AF software (Leica Microsystem, Inc.) and Imaris software v.9.9.1 64× (Oxford Instruments plc). The intensities of fluorescent signals in regions of interests (ROI) were measured by LSA AF Lite software (Leica Microsystem).

### *In vitro* imaging of HEK293 cells

HEK293 cells were transfected using TransIT-293 transfection reagent (Mirus Bio LLC). For transient expression of human Siglec-5-GFP 80% confluent HEK293 in 8 well chamber slide (0.25 ml/well) were transfected by adding dropwise to each well 25 μl containing 0.25 μg of plasmid diluted in Opti-MEM with a 1:4 (DNA/reagent) dilution of TransIT-293 Transfection Reagent. To generate a stable cell line of human Siglec-5-GFP and human ACE2-OFP transfected HEK293 cells were sorted with FACS Aria II by GFP and OFP signals. Sorted human Siglec-5-GFP transfected HEK293 cells were further selected with G418 (0.8 mg/ml) containing growth media and maintained with G418 (0.5 mg/ml) media. Sorted human ACE2OFP transfected HEK293 cells were further selected with hygromycin B (0.2 mg/ml) containing growth media and maintained with hygromycin B (0.1 mg/ml) media. Transiently transfected HEK293 cells in 8 well chamber slide were directly imaged with a Leica SP8 inverted five channels confocal microscope (Leica Microsystems). HEK293 cells, HEK293 Siglec-5-GFP, and HEK293 ACE2-OFP cell lines were plated at a ratio of 1:1:1 (cell numbers) in 8 well chamber with normal media 18 hr prior to imaging. Imaging was performed with a confocal microscope equipped with 40 × oil objective, 0.95 NA (immersion medium used distilled water). The temperature of air (5% CO_2_) was maintained at 37.0 ± 0.5°C. Fluorescent SARS-CoV-2 Spike protein (Alexa Fluor 647) (1 μg/ml) was added into culture media. Images were acquired with Leica LAS AF software (Leica Microsystem, Inc) and processed with Imaris software v.9.9.1 64× (Oxford Instruments plc).

### Time-lapse imaging of lung sections with confocal microscopy

Lung slices were obtained from mouse lungs using a slightly modified published protocol (53). Briefly, C57BL/6 mice were euthanized with overdose of Avertin (1 ml of 2.5% Avertin). The peritoneum was opened, and the descending aorta cut allowing blood to pool in the abdomen. The trachea was cannulated, and the lungs inflated with 37 °C 1.5% low-melting-point agarose (Cat# 50111, Lonza) prepared with RPMI 1640 media. Subsequently, the lungs were excised and rinsed with RPMI 1640 media. Isolated left lungs were sectioned into six to eight 1-mm-thick transverse slices using a #10 scalpel blade. Lung slices were placed in a pre-warmed cover glass chamber slide (Nalgene, Nunc) under a metal flat washer (M8-5/16^th^ inches diameter). The chamber slide was then placed into the temperature control chamber on the microscope. The temperature of air was monitored and maintained at 37.0± 0.5°C for 5% CO_2_. Mounted lung sections on the chamber slide were microscopically analyzed using a Leica SP8 confocal microscope equipped with an HC PL APO CS2 40× (NA, 1.30) oil objective (Leica Microsystem, Inc.) and images were processed with Leica LAS AF software (Leica Microsystem, Inc.). Lung slices were imaged a range of depths (10-50 μm). Time-lapse images were processed with Imaris software v.9.9.1 64× (Oxford Instruments plc).

### Intravital imaging

The microanatomy of liver, spleen, heart, Peyer’s patch, and inguinal LN were delineated by tail vein injection of labelled antibodies before imaging. Antibodies used included CD31, blood vessels; F4/80, Kupffer cells and macrophages; CD169, subcapsular macrophages; and Ly6G; neutrophils). The antibody mixtures were injected 10 min before starting animal surgery. Fluorescently labelled Spike proteins were injected intravenously or at the mouse tail base as indicated. To image the liver or spleen a slightly modified published protocol was used (54). Briefly, after initial anesthesia (Avertin 300 mg/kg, i.p.) the skin and peritoneum were cut to expose the left lobe of the liver, or the left flank was cut below the costal margin to expose the spleen. The visible organs were glued with n-butyl cyanoacrylate to a custom-made metal holder. After attachment, the mouse was placed over a pre-warmed cover glass (Brain Research Laboratories) window on universal mounting frame AK-Set (PECON^®^). The exposed organs were kept moist with saline wetted gauze. The mounting frame was placed into the temperature control chamber on the microscope and maintained at 37.0± 0.5°C. Once stabilized onto imaging stage/insert the mice received isoflurane (Baxter; 2% for induction of anesthesia, and 1 – 1.5% for maintenance, vaporized in an 80:20 mixture of oxygen and air). For liver and spleen four-dimensional analysis of cell behavior, stacks of various numbers of section (z step = 3, 5) were acquired every 5-30 sec to provide an imaging volume of 20 ∼ 50 μm in depth. Intact organ imaging of the heart, inguinal lymph node, or Peyer’s patches was performed after mouse sacrifice using same imaging procedure as used for liver/spleen imaging.

To image neutrophils in the cremaster muscle (31), mice received an intrascrotal injection of PBS, BSA, IL-1β (50 ng in 300 µl saline, R&D Systems), Spike proteins, or IL-1b and Spike proteins. 90 min. prior to imaging, the mice received injections of Avertin (300 mg/kg, i.p.) and fluorescently labeled antibodies intravenously. Antibodies used directed against Gr-1, neutrophils; GPIbβ, platelets; and CD31, blood vessel endothelium. The isolated cremaster tissue was exteriorized and stabilized onto the imaging stage/insert with the tissue directly contacting the cover glass. The exposed tissue was kept moist with prewarmed saline (37°C). Once stabilized onto an imaging stage/insert the mouse received isoflurane (Baxter; 2% for induction of anesthesia, and 1 – 1.5% for maintenance, vaporized in an 80:20 mixture of oxygen and air), and placed into a temperature-controlled chamber. Image stacks of optical sections (z step=1) were routinely acquired at 20 – 30 sec intervals to provide an imaging volume of 15-25 µm. All imaging was performed with a Leica SP8 inverted 5 channels confocal microscope (Leica Microsystems) equipped with HC PL APO CS2 40× (NA, 1.30) oil objective. Sequences of image stacks were transformed into volume-rendered four-dimensional videos using Imaris software v.9.9.1 64x (Oxford Instruments plc). Video editing was performed using Adobe Premiere Pro 2022 (Adobe Systems Incorporated).

### Quantifications and Statistical Analyses

All experiments were performed at least three times. Representative images were placed in figures. Primary image data which was analyzed and calculated by Leica LAS AF software (Leica Microsystem, Inc.) or Imaris software v.9.9.1 64× (Oxford Instruments plc). was acquired and processed with Microsoft Excel software. Error bars with ±SEM, and p values were calculated with unpaired t-test in GraphPad Prism 9.3.1 (GraphPad software). p<0.05 was considered significantly different.

## Acknowledgements

The authors thank Dr. Anthony Fauci for long standing encouragement.

## Author Contribution

CP performed most of the mouse work; IYH performed the flow cytometry and macrophage cytokine production; SLY performed the cremaster imaging, SV, DW, and DVR purified the recombinant proteins, CC and JA supervised the protein purification and provided helpful advice, JHK wrote the manuscript and oversaw the project.

## Data and Material Availability

The primary imaging files are available from the corresponding author.

## Supplementary Material

**Figure 1-figure supplement 1.**
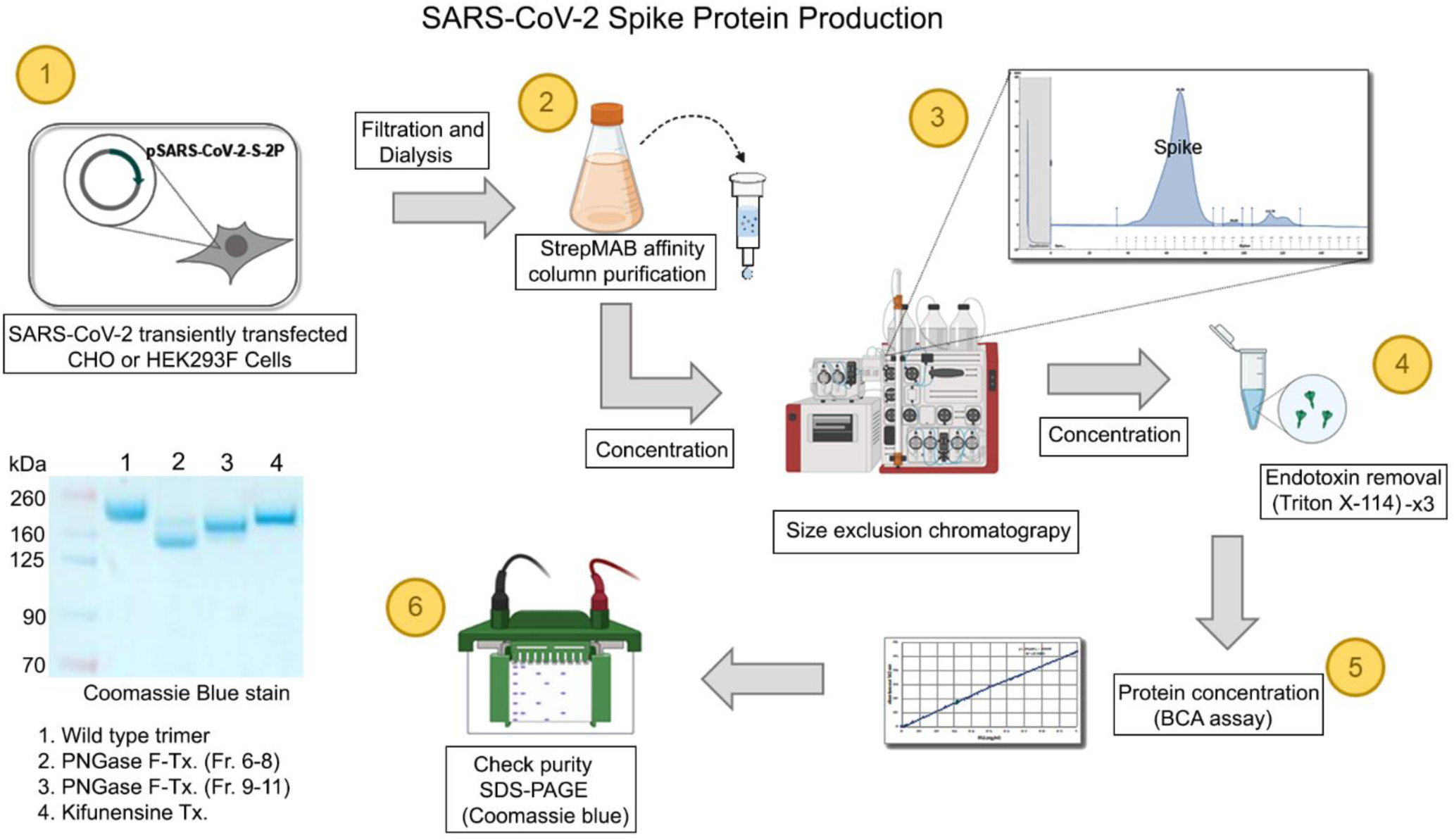
Preparation of SARS-CoV-2 proteins. Outline of recombinant proteins production and purification from cultured media of pSARS-CoV-2-S-2P transfected CHO or HEK293F cells. See methods for PNGase F and Kifunensine treatment. Calculated molecular weights: wild type trimer −210 kDa, PNGase F-Tx. (Fr. 6-8)-146 kDa, PNGase F-Tx (Fx. 9-11)-178 kDa, and Kifunensine Tx.-197.5 kDa.

**Figure 1-figure supplement 2.**
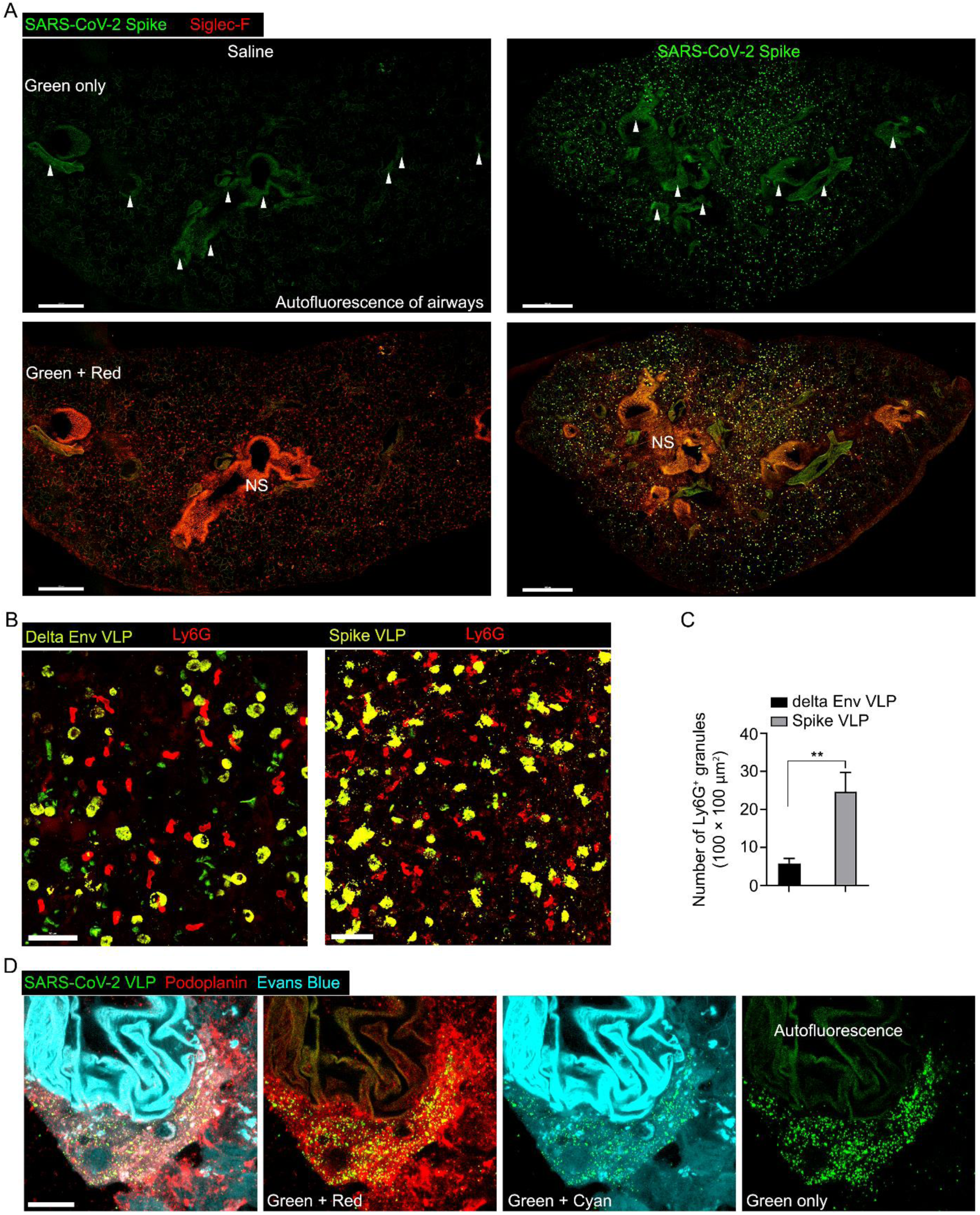
Confocal micrographs were taken for staining control and to analyze neutrophil fragmentation caused by SARS-CoV-1 Spike VLPs, comparing them to delta Env VLPs. **(A)** Confocal micrographs compare the negative control (saline) with the positive sample (SARS-CoV-2 Spike, Alexa Fluor 488). Arrowheads in the upper panels indicate autofluorescence background. The nonspecific Siglec-F antibody stain background is denoted as ‘NS’ in the lower panels. Scale bars, 500 μm. **(B)** Confocal micrographs of lungs collected at 3 hours post instillation of SARS-CoV-2 Spike protein (left) or delta envelope (right) VLP (green, GFP) are shown. Infiltrated neutrophils (red, Ly6G) in lung tissue were visualized. Scale bars, 50 μm. (**C**) Neutrophil fragmentation was measured by counting the Ly6G+ granules in five different areas of 100 × 100 μm² each. ** p < 0.01, paired t-test. (**D)** A confocal micrograph shows a lung lymphatic vasculature visualized with Podoplanin antibody. Fifty microliters of a mixture of Evans blue (cyan) (5 μg) and Spike bearing VLPs (green) (0.5 million counts) were applied to the mouse nose. The first panel presents the merged image of all signals, while the subsequent panels display specific color combinations: green and red in the second, and green and cyan in the third. The fourth panel exclusively exhibits the green signal, with autofluorescence of the lung airway structure indicated. Scale bars, 20 μm.

**Figure 2-figure supplement 1.**
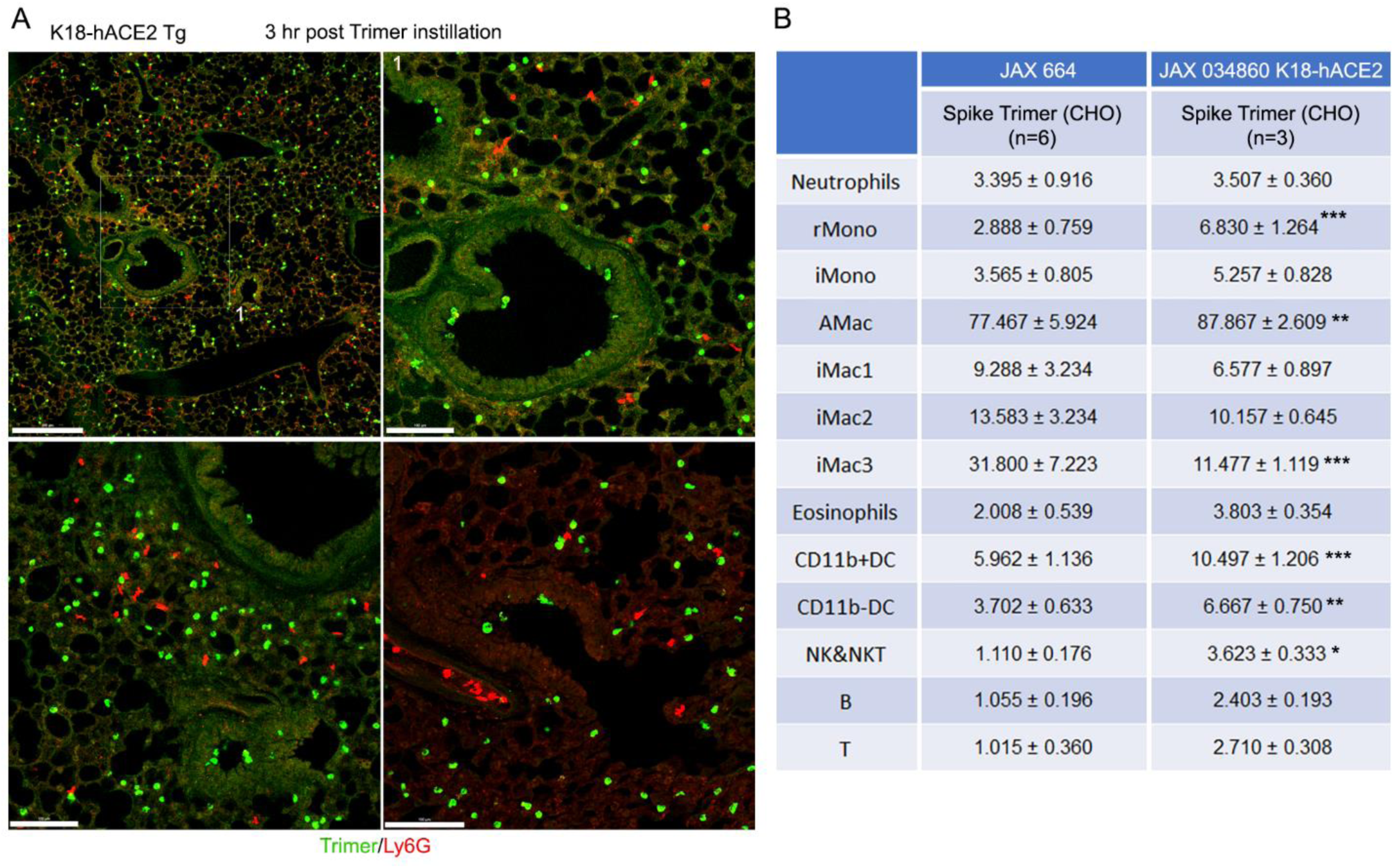
Intranasal SARS-CoV-2 administration to K18-hACE2 mice. (**A**) Representative confocal images of lung sections 3h post intranasal SARS-CoV-2 Spike protein (3 µg, green). Neutrophils immunostained with Ly6G. (**B**) SARS-CoV-2 Spike protein uptake by lung leukocytes 18 hr following intranasal inoculation. Flow cytometry results from analysis of leukocytes purified from the lungs of WT or K18-hACE2 mice. Data for JAX 664 mice same as shown in figure 2. K18-hACE2 leukocyte values significantly different from those from the JAX 664 mice are indicated, *p < 0.05; **p < 0.01; ***p < 0.001.

**Figure 3-figure supplement 1.**
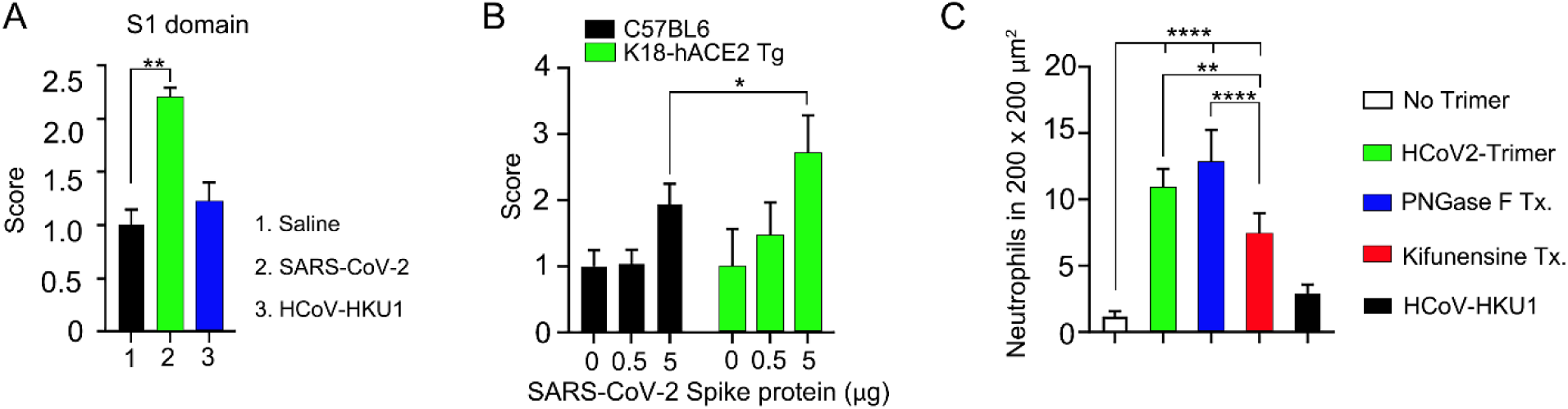
Quantification of lung permeability and neutrophil recruitment. **(A)** Evans blue (200 µl of 5 mg/ml in PBS) was intravenously injected 1.5 hours after intranasal Spike administration. The indicated recombinant proteins’ S1 domains (3 μg per mouse) were administered. Lungs were collected 1 hour after Evans blue injection. The score was calculated based on the amount of Evans blue dye (in μg/g) in the lungs administered with the Spike protein, divided by the average amount of Evans blue dye in lungs administered with saline. ** p < 0.01, One-way ANOVA. **(B)** The permeability of lung vasculature was measured by comparing wild-type mice with K18-hACE2 transgenic mice. The SARS-CoV-2 Spike protein was administered at two different concentrations: 0.5 μg or 5 μg per mouse. * p < 0.05, Two-way ANOVA. **(C)** The number of neutrophils was counted in six different areas, each measuring 200 × 200 μm². ****p < 0.0001, One-way ANOVA.

**Figure 4-figure supplement 1.**
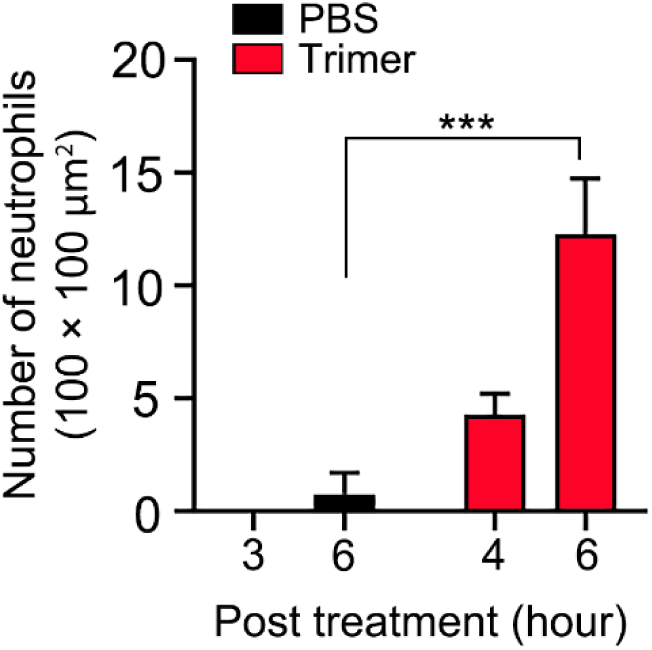
Quantification of neutrophil count in cremaster muscle. The number of neutrophils was counted in four 100 × 100 μm² areas. ***p < 0.001, Two-way ANOVA.

**Figure 4-figure supplement 2.**
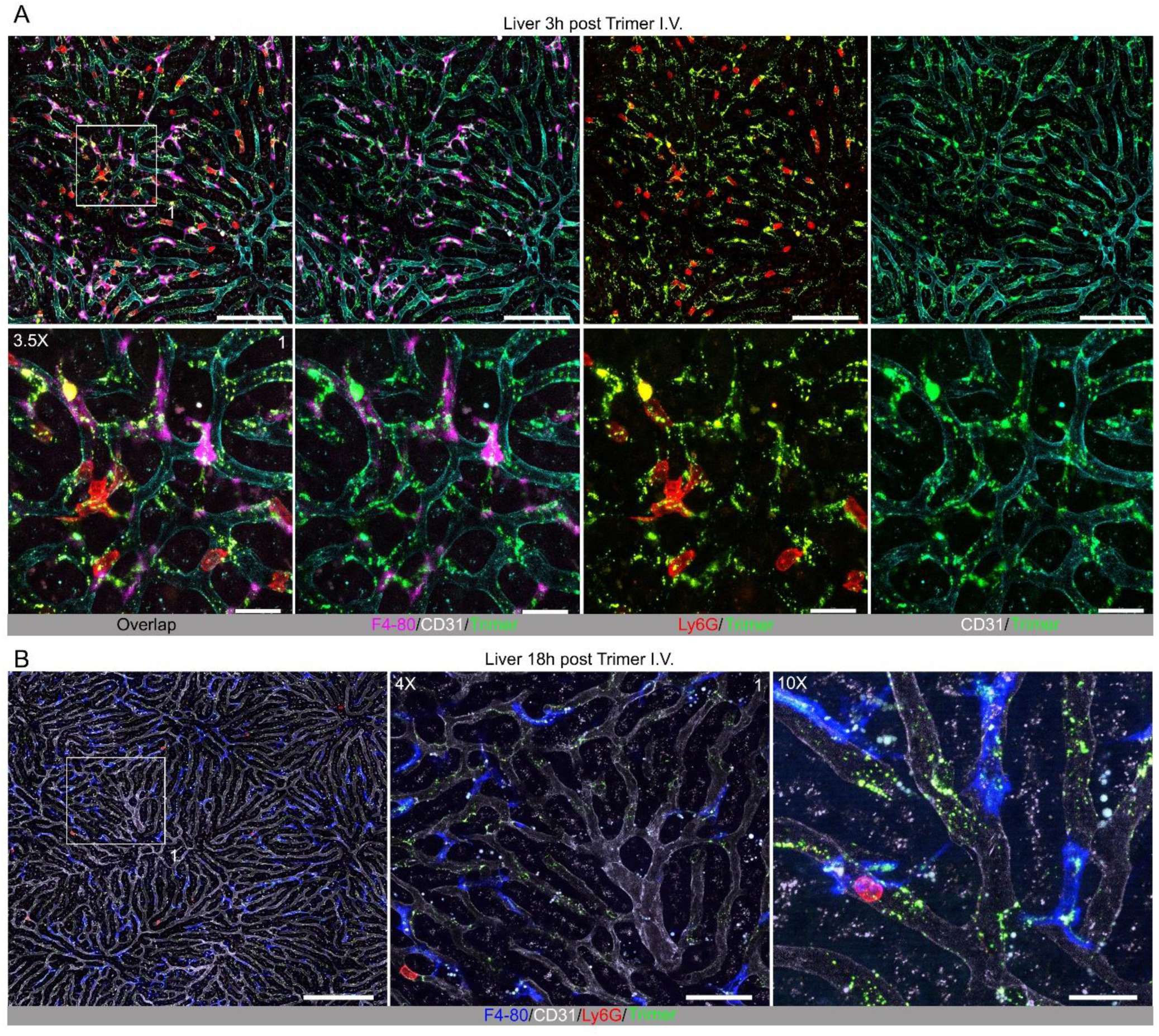
Localization of Spike protein in the liver following intravenous administration. **(A)** Confocal micrographs of the liver imaged at 3 hr post intravenous injection of SARS-CoV-2 Spike (Trimer) protein. Spike protein (green), Kupffer cells (magenta, F4/80), and neutrophils (red, Ly6G) shown in liver sinusoid vasculature (cyan, CD31). Antibodies injected 0.3 hr before imaging. ROI-1 (box) (upper left) is enlarged (3.5× magnification) in lower panels. Scale bars, 100 and 20 μm. (**B)** Confocal micrographs of liver imaged at 18 hr post intravenous injection of SARS-CoV-2 Spike protein. Spike protein (green), Kupffer cells (blue, F4/80), and neutrophils (red, Ly6G) are shown in liver sinusoid vasculature (white, CD31). Antibodies injected 0.3 hr before imaging. ROI-1 (box) (left) is enlarged in the middle panel (4× magnification). Enlarged image shows neutrophil contacting Spike protein bearing cells on liver sinusoid endothelium (10× magnification). Scale bars, 200, 50, and 20 μm.

**Figure 4-figure supplement 3.**
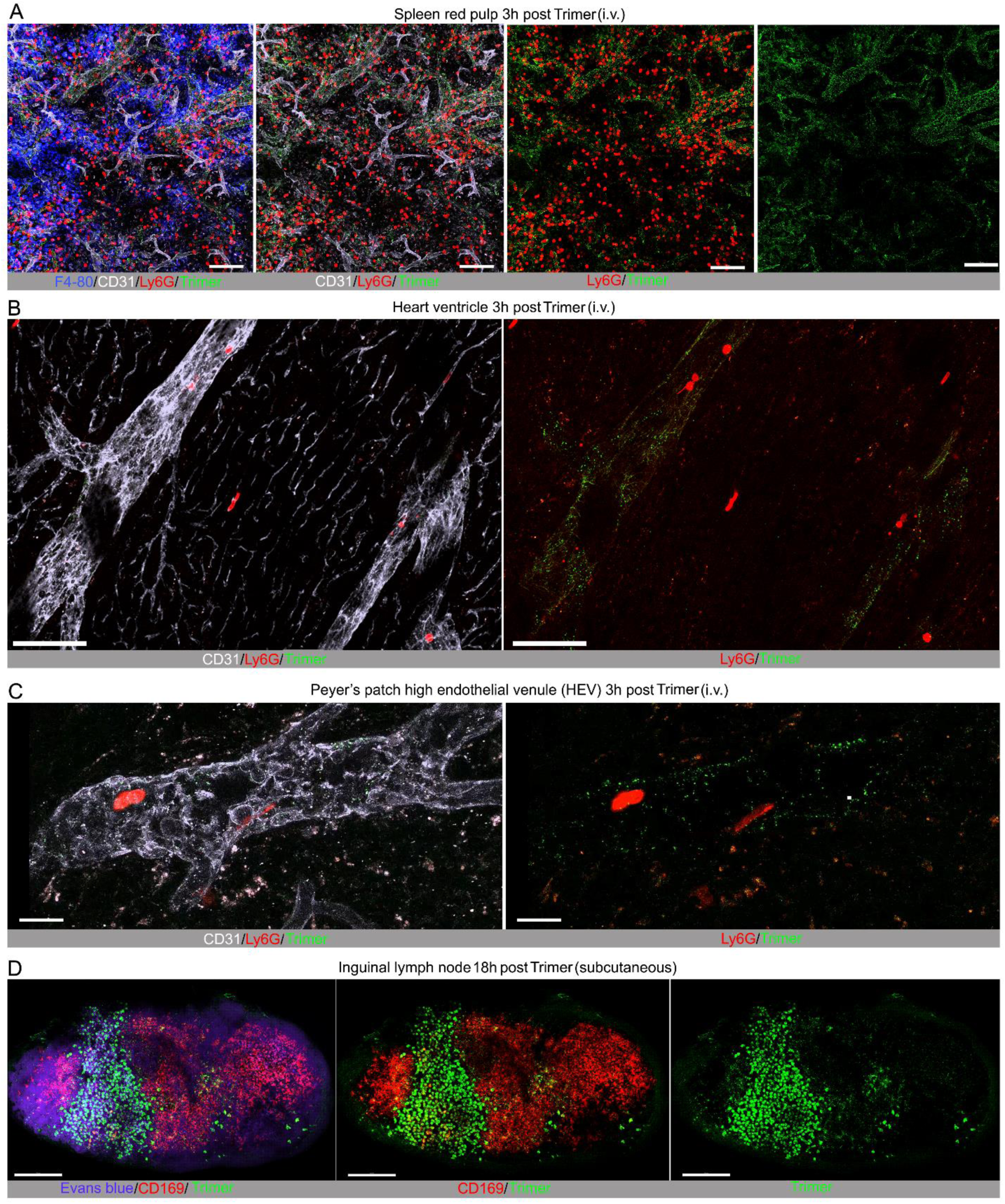
Localization of SARS-CoV-2 Spike protein at other sites following intravenous injection and subcutaneous injection near the inguinal lymph node. **(A)** Confocal micrographs of mouse spleen imaged 3 hr post intravenous injection of SARS-CoV-2 Spike (Trimer) protein. Spike protein (green), red pulp macrophages (blue, F4/80), and neutrophils (red, Ly6G) visualized in spleen red pulp vasculature (cyan, CD31). Antibodies injected 1 hr before imaging. Scale bars, 100 μm. (**B)** Confocal micrographs of heart ventricle imaged 3 hr post intravenous injection of Spike protein. Spike protein (green) and neutrophils (red, Ly6G) localized in heart blood vessels (white, CD31). Antibodies injected 1 hr before imaging. Scale bars, 100 μm. (**C)** Confocal micrographs of Peyer’s patch high endothelial venules (HEV) imaged at 3 hr post iv injection of Spike protein. Spike protein (green) and neutrophils (red, Ly6G) shown in HEV (white, CD31). Antibodies injected 0.5 hr before imaging. Scale bars, 20 μm. (**D)** Confocal micrographs of inguinal lymph node imaged at 18 hr post tail base injection of Spike protein. Spike protein (green) and subscapular sinus macrophages (red, CD169) shown. Subcapsular sinus defined via Evans blue (purple) signal. Antibodies and Evans blue injected 1 hr before imaging. Scale bars, 200 μm.

## Video Legends

**Video 1.** Local Spike protein injection results in neutrophil recruitment in the cremaster muscle. Confocal intravital microscopy movie shows neutrophils (green), platelets (red), endothelium (blue) immunostained with Alexa Fluor 488-Gr1, DyLight 649-GP1bβ, and Alexa Fluor 555-CD31 monoclonal antibodies, respectively. Intrascrotal injection of PBS or Spike protein (5 µg) 3-4 hours prior to imaging. The first sequence spans a ∼60 min (PBS) while the second sequence spans a ∼45 min (Spike) imaging period. Images were captured at ∼1 frame per 30s with ∼100× magnification. Time counter is hour:minute:second.

**Video 2.** Local injection of Spike proteins causes neutrophil fragmentation. Confocal intravital microscopy movie shows neutrophils (green), platelets (red), endothelium (blue) immunostained with Alexa Fluor 488Gr1, Dylight 649-GP1bβ, and Alexa Fluor 555-CD31 monoclonal antibodies, respectively. Intrascrotal injection of Il-1β or Il-1β plus 5 µg Spike protein ∼20 hours prior to imaging. Both sequences span a 20 min period. Images were captured at ∼1 frame per 30s with ∼100× magnification. Time counter is hour:minute:second.

**Video 3.** Intravenously injected Spike protein outlines liver sinusoids and accumulates on Kupffer cells. The liver imaging was taken at three different time points; from 20 min to 1hr, from 1hr 10 min to 1hr 15 min, and from 1hr 25 min to 46 min after SARS-CoV-2 Spike protein (green, Alexa Fluor 488) injection. An image sequence of a 12 μm z-projection was acquired with 40× lens as scanning speed of 11.05 sec between frames. Kupffer cells (magenta, F4/80), and neutrophils (red, Ly6G) in liver sinusoid vasculature (cyan, CD31) were visualized with antibody injection into tail vein at 30 min prior to imaging. The scale bar represents 20 μm. Time counter is hour:minute:second.

**Video 4.** Neutrophils undergo NETosis when plated on A549 cells in the presence of Spike protein. A549 cells were plated on chamber slide 48 hr before trimer treatment. SARS-CoV-2 Spike trimer (green) was added to A549 cell culture 24 hr before neutrophil seeding. Purified human neutrophils stained with Hoechst (cyan) and were seeded on A549 cells in propidium iodide (PI, 1 μg/ml) containing culture media. NETosis of neutrophils were detected by exposed DNA (red, PI). An image sequence of a 20 μm zprojection was acquired with 40× lens at a scanning speed 1 frame /10 min over 5 hours. Regions of interest (Box A and B) demonstrate a typical NETosis of neutrophil contacting on trimer bearing A549 cell were enlarged in second part of video. Scale bars, 30 and 10 μm. Time banner, hour:minute:second.

**Video 5.** hSiglec-5 expressing HEK293 cells bind and endocytosis Spike protein. Individually established stable-transfected cells expressing Siglec-5-GFP or ACE2-OFP were plated in the same chamber slide with non-transfection cells. After overnight culture SARS-CoV-2 Spike protein (1 μg/ml) was overlaid for 1 hr before imaging. An image sequence of a 3 μm z-projection was acquired with 40× lens at a scanning speed of 23 sec between frames. Signals visualized as SARS-CoV-2 Spike trimer (blue), Siglec-5-GFP (green), ACE2-OFP (red), and non-transfected HEK293 cells (gray). The scale bar represents 20 μm. Time counter is hour:minute:second.

---

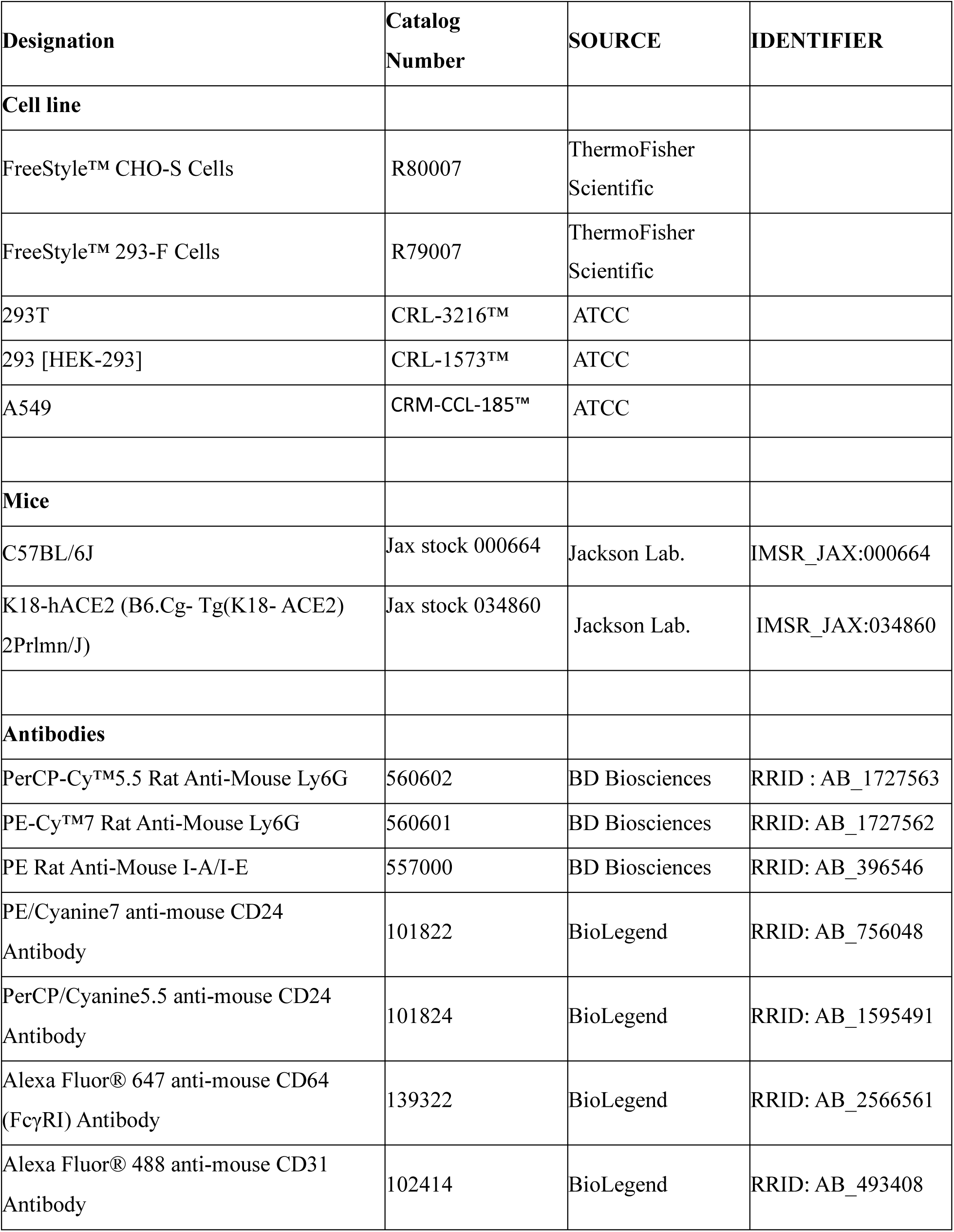
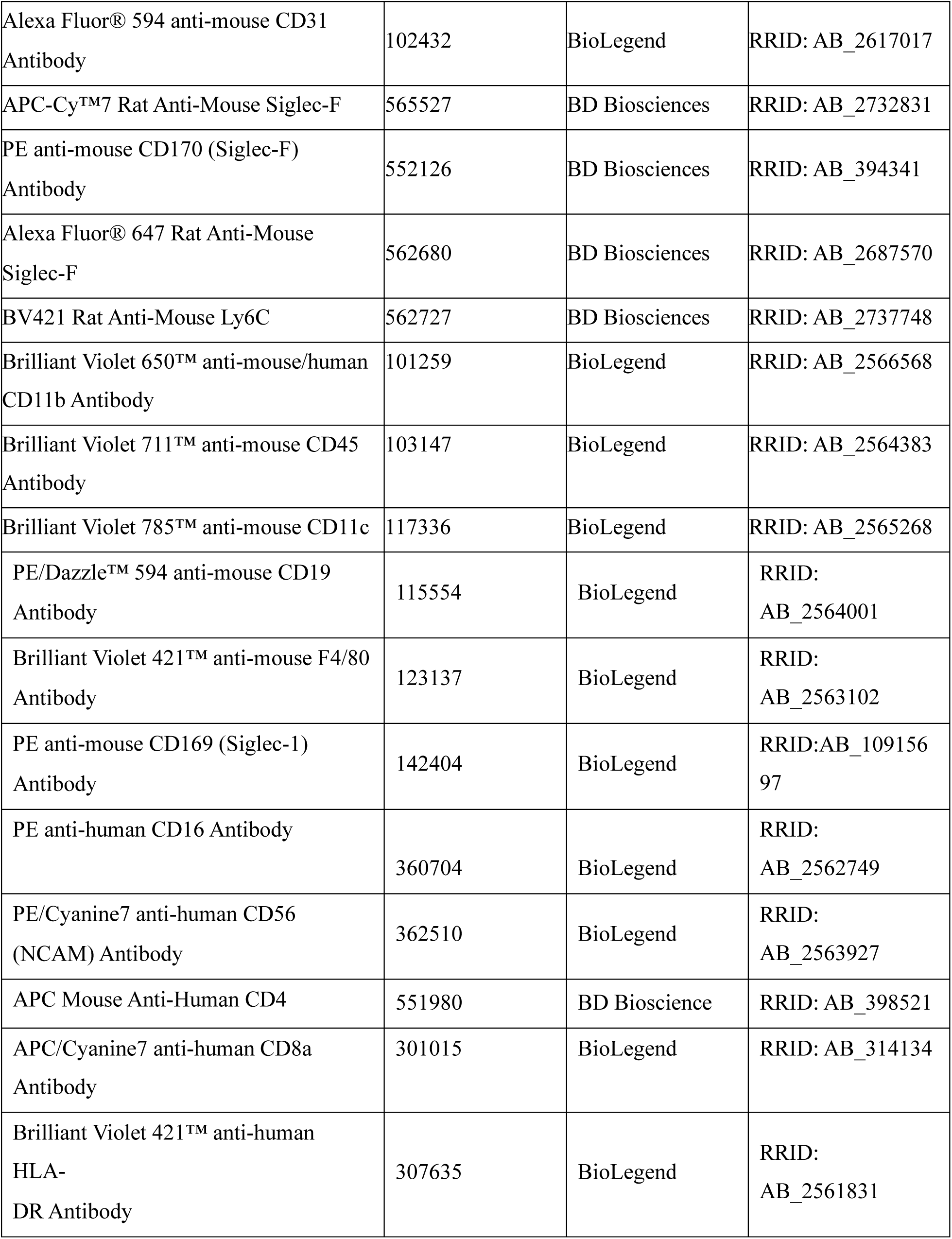
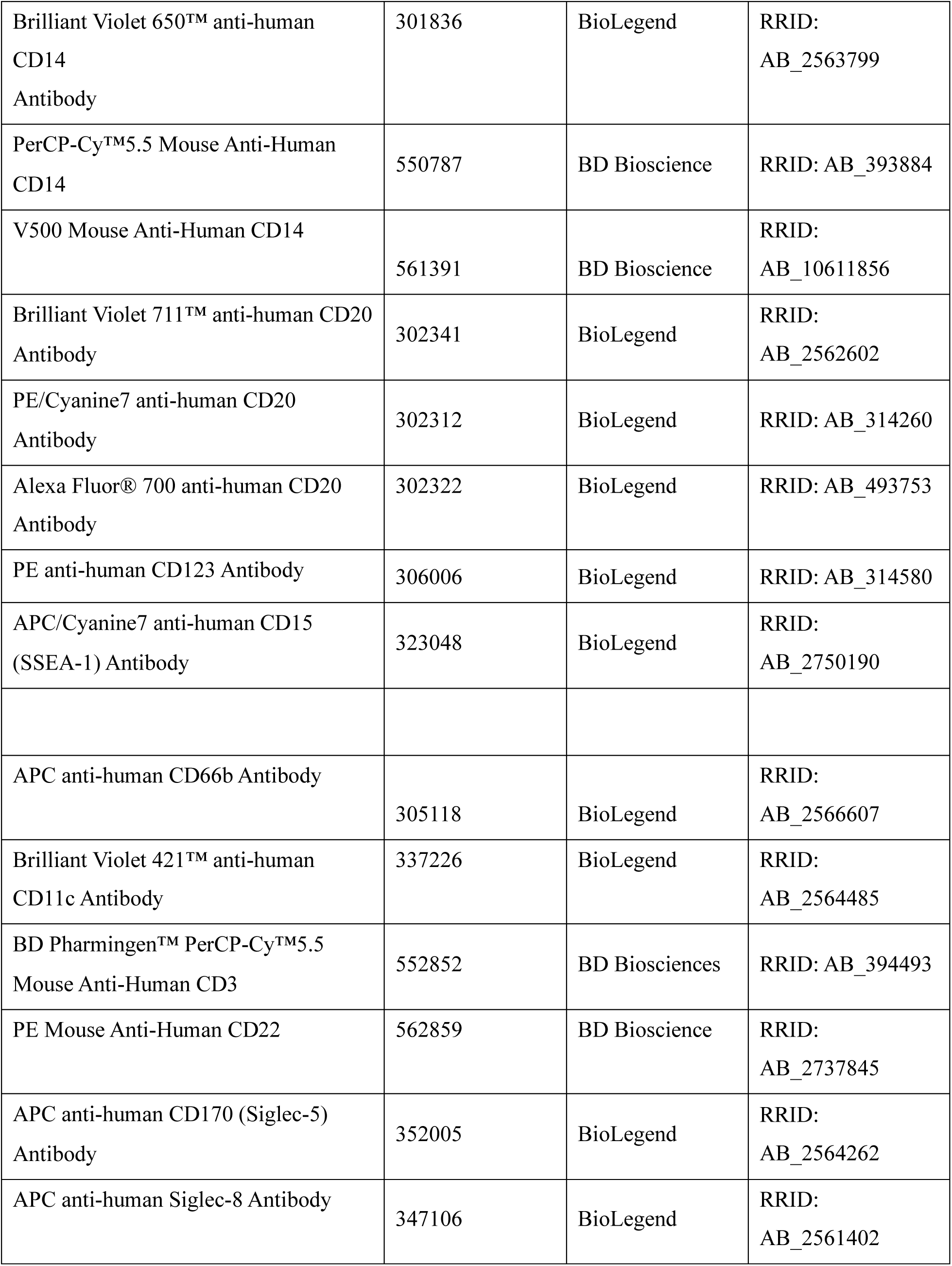
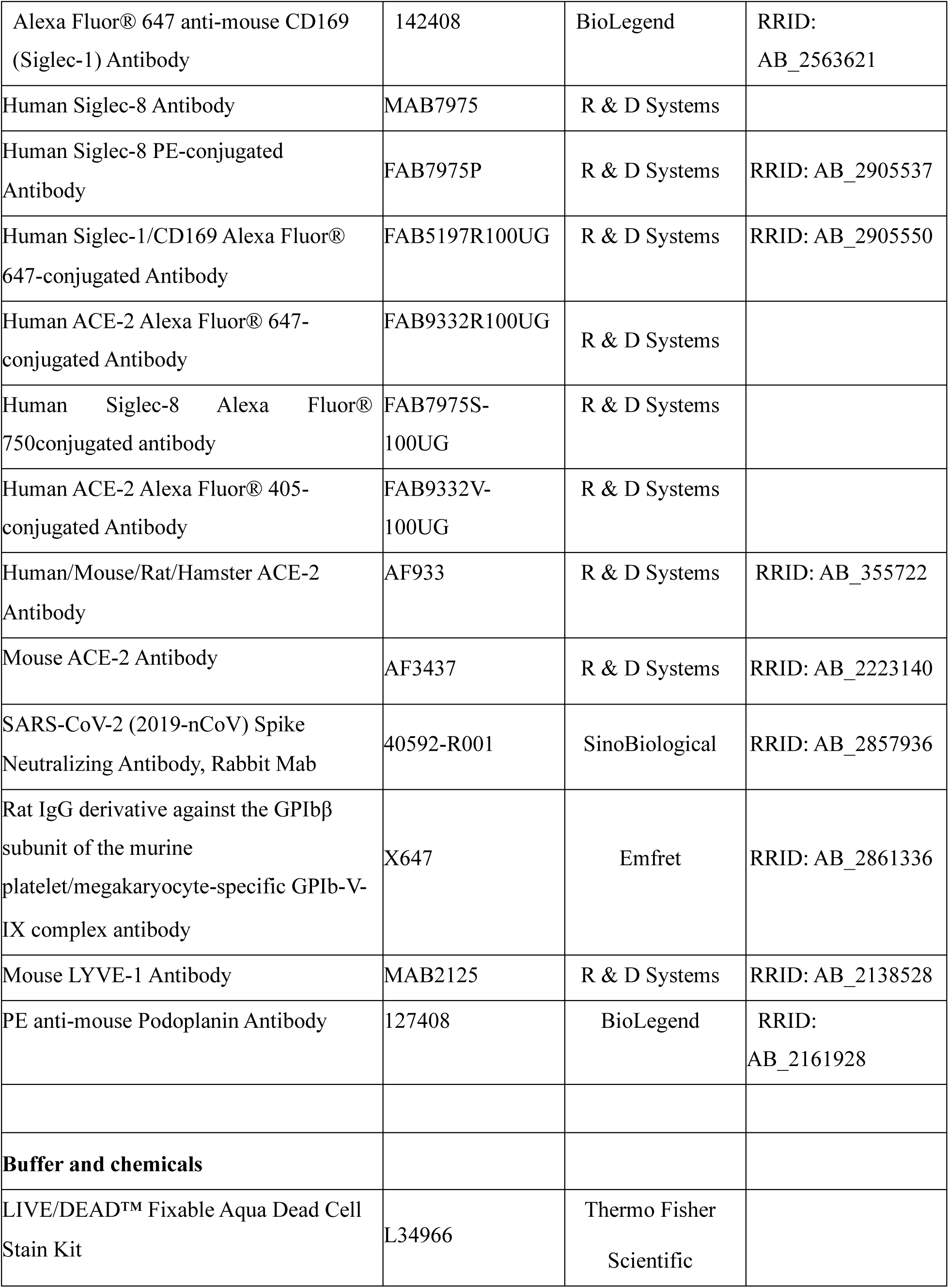
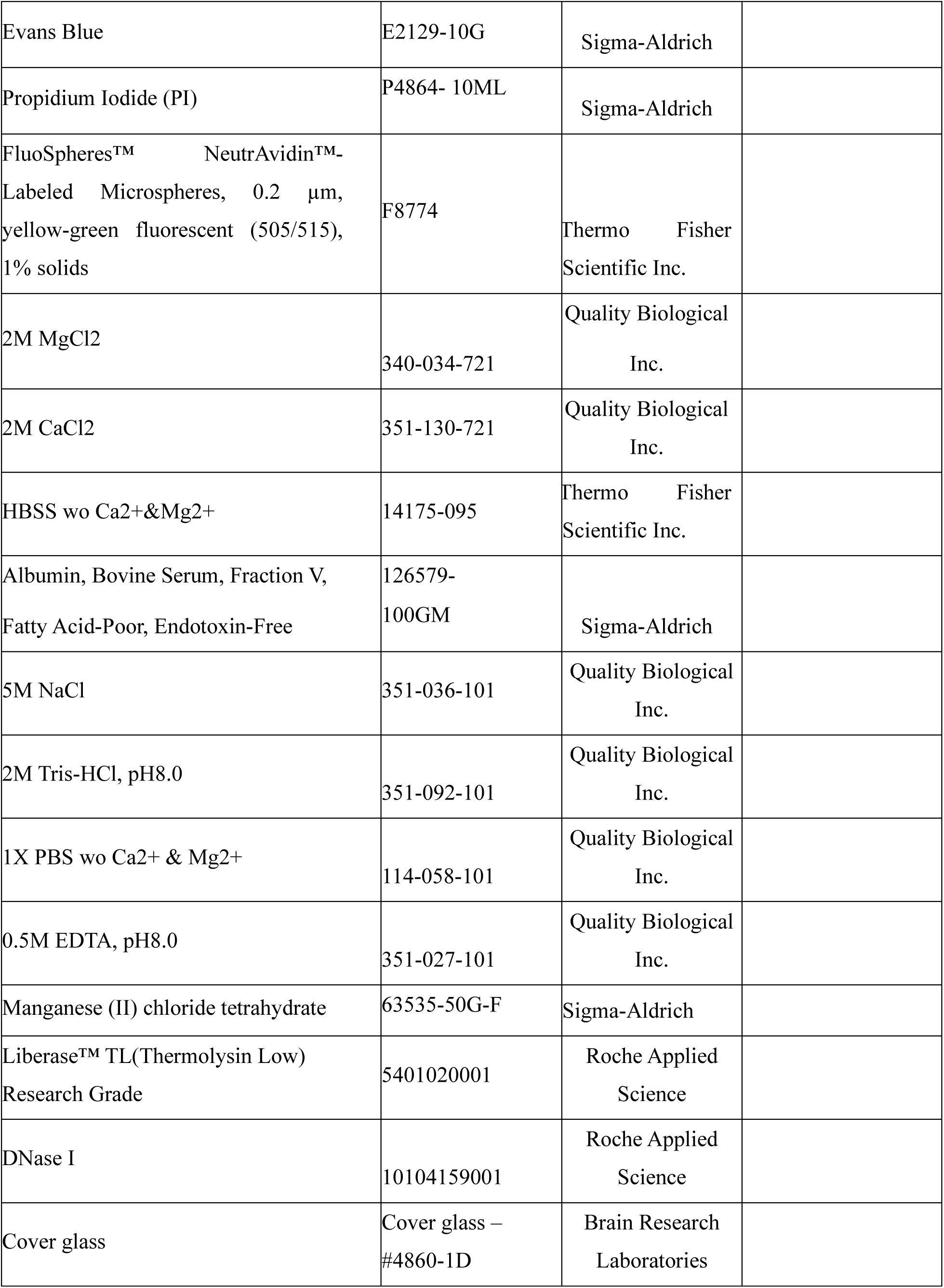
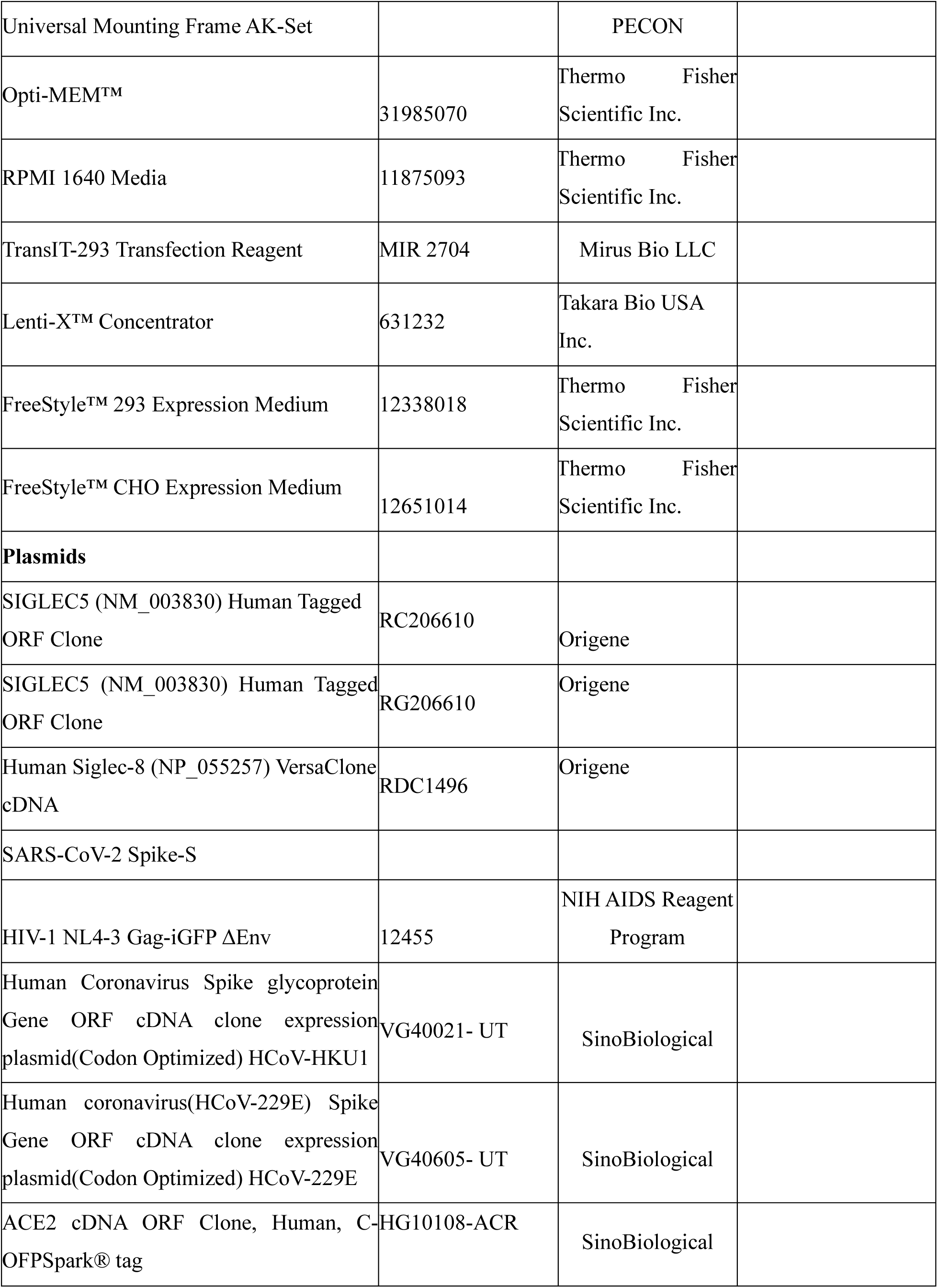
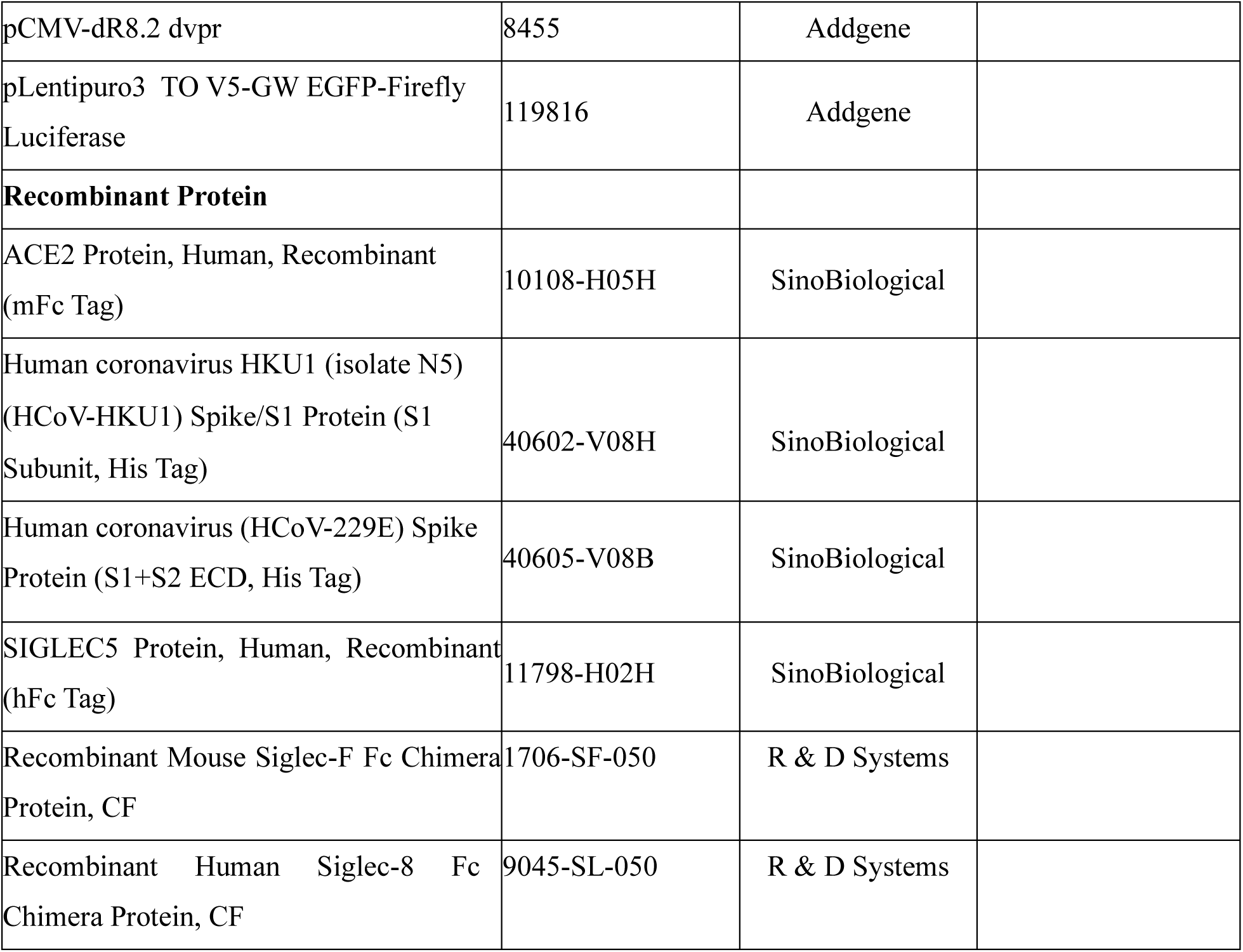
Reagent Table.

